# Learning reorganizes dendritic and stabilizes axonal initial segment inhibitory synapses in CA1 pyramidal neurons

**DOI:** 10.1101/2025.03.04.641429

**Authors:** H. Klimmt, D. Kappel, A.F. Ulivi, A.Ö. Argunsah, B. Murthy, S. Somatakis, R.E. Huettl, S. Remy, A. Attardo

**Affiliations:** Leibniz Institute for Neurobiology, Magdeburg, Germany; International Max Plank Research School for Translational Psychiatry, Munich, Germany; Center for Cognitive Interaction Technology, Bielefeld University, Bielefeld, Germany; Brain Research Institute (HiFo), University of Zurich and Neuroscience Center Zurich, Zurich, Switzerland; Graduate School for Systemic Neuroscience, Ludwig-Maximilian University, Munich, Germany; Max Planck Institute of Psychiatry, Munich, Germany; German Center for Mental Health (DZPG), partner site Halle-Jena-Magdeburg and Center for Behavioral Brain Science (CBBS), Magdeburg, Germany; German Center for Neurodegenerative Diseases (DZNE), Magdeburg, Germany

**Keywords:** Inhibitory synapses, Structural plasticity, Learning, Deep-brain two-photon optical imaging, Mathematical modelling, Axon initial segment

## Abstract

Structural synaptic plasticity underlies the changes in brain connectivity required for learning and memory. Inhibitory synapses (INS) target all subcellular domains of excitatory pyramidal neurons (PNs), including dendrites, somata and axon initial segments (AIS). These subcellular domains have distinct molecular, structural and physiological profiles which underlie their functions. How structural plasticity of INS supports these functions as well as emerging properties such as memory is largely unknown. To tackle these questions we tracked INS on dendrites, somata and AIS of PNs in the dorsal hippocampal CA1 area of mice over two weeks. Size and temporal dynamics of INS showed a strong compartmentalization and dendritic INS were less dynamic than dendritic spines. Trace fear conditioning led to reorganization of dendritic INS and to stabilization of AIS INS but had a minimal effect on dendritic spines. Finally, mathematical modelling allowed us to probe the mechanisms underlying stabilization of INS upon learning.

## INTRODUCTION

Coordinated changes in synaptic connections give rise to the brain’s ability to learn and recall information^1–3^. These modifications include strengthening or weakening of existing synapses, as well as synapse formation and elimination. Structural synaptic plasticity underlies long-term changes in connectivity, such as the ones that might be required for learning and memory ^4,5^. Moreover, the interplay between excitatory and inhibitory synaptic transmission serves an important role in adult brain and plasticity of excitatory and inhibitory inputs both participate in the processing and integration of local dendritic activity ^6,7^.

Inhibitory synapses (INS) target all subcellular domains of excitatory pyramidal neurons (PNs), including the dendritic arborizations, the soma and the axon initial segment (AIS) ^8,9^ . These subcellular domains have distinct molecular, structural and physiological profiles which underlie their specific computational functions: dendrites receive and compute all excitatory inputs, while somata and AIS provide the cellular output. The function of the different classes of inhibitory neurons projecting to these subcellular compartments has been the subject of intense investigation ^10–12^ , yet how the structural plasticity of the INS supports these functions is largely unknown. Likewise, different classes of inhibitory neurons projecting to different subcellular compartments have been implicated in different aspects of hippocampal learning ^13–18^, but if and how structural INS plasticity in these compartments relate to learning has not been established.

Longitudinal two-photon (2P) optical imaging enabled investigating long-term dynamics of excitatory synaptic structural plasticity in the brain of live animals and led to a leap in understanding of brain plasticity and its role in learning and recall ^19–21^. By comparison the structural plasticity of INS is understudied. Moreover, the vast majority of previous work mostly focused on neocortical areas ^22–27^, thus leaving other brain areas key for learning and memory – such as the hippocampus – unexplored.

Here we confronted this knowledge gap by labelling INS impinging on PNs located in the dorsal hippocampal CA1 area (dCA1) of mice and by tracking them over two weeks by using deep-brain 2P longitudinal optical imaging. This allowed us to simultaneously monitor INS located on basal dendritic arborizations, somata and AIS of dCA1 PNs and to characterize the long-term structural dynamics of these INS. Then, we compared dynamics of inhibitory and excitatory synapses - by using dendritic spines as a proxy - during baseline conditions and upon learning of a hippocampal-dependent task. Size and temporal dynamics of INS showed a very strong compartmentalization with soma INS being the most dynamic and dendritic INS were overall less dynamic than dendritic spines. Trace fear conditioning led to highly compartmentalized changes in INS long-term temporal dynamics but not in dendritic spines. Finally, we used mathematical modelling to investigate the circuit mechanisms underlying stabilization of AIS INS upon learning of a hippocampal-dependent task.

## RESULTS

### Longitudinal tracking of INS in different subcellular compartments of dCA1 PNs

We adapted a method previously developed to genetically tag INS in the brain of live mice ^22,24^ and labelled the morphology of PNs and the INS impinging on them. We transduced the dCA1 of C57Bl6 mice with three Adeno Associated Viruses encoding for (i) a Cre-recombinase under the control of the pan-excitatory neuron promoter CaMKII alpha, (ii) a Cre-dependent, cell-filling, red fluorescent protein (tdTomato) and (iii) a Cre-dependent, green fluorescent protein (GFP) fused to the GABAergic postsynaptic protein Gephyrin (**Fig. 1a**). We thus detected INS on dendrites, somata and AIS of a subset of dCA1 PNs. AIS INS were identified by their large size - consistent to cartridge synapses ^28–32–^ and confirmed by AnkyrinG immunostaining (**Fig. 1b**). We used longitudinal 2P optical imaging to image INS impinging on these subcellular compartments. (**Fig. 1c**). We tracked dendritic, somatic and AIS INS (**Fig. S1a**) for a duration of 13 days. In total, we sampled 507 dendritic, 472 somatic and 169 AIS independent inhibitory postsynaptic sites, *i.e.* the positions on a PN where we could detect an INS at least once during the experiment (**Fig. S1b-c**).

**Figure 1.**
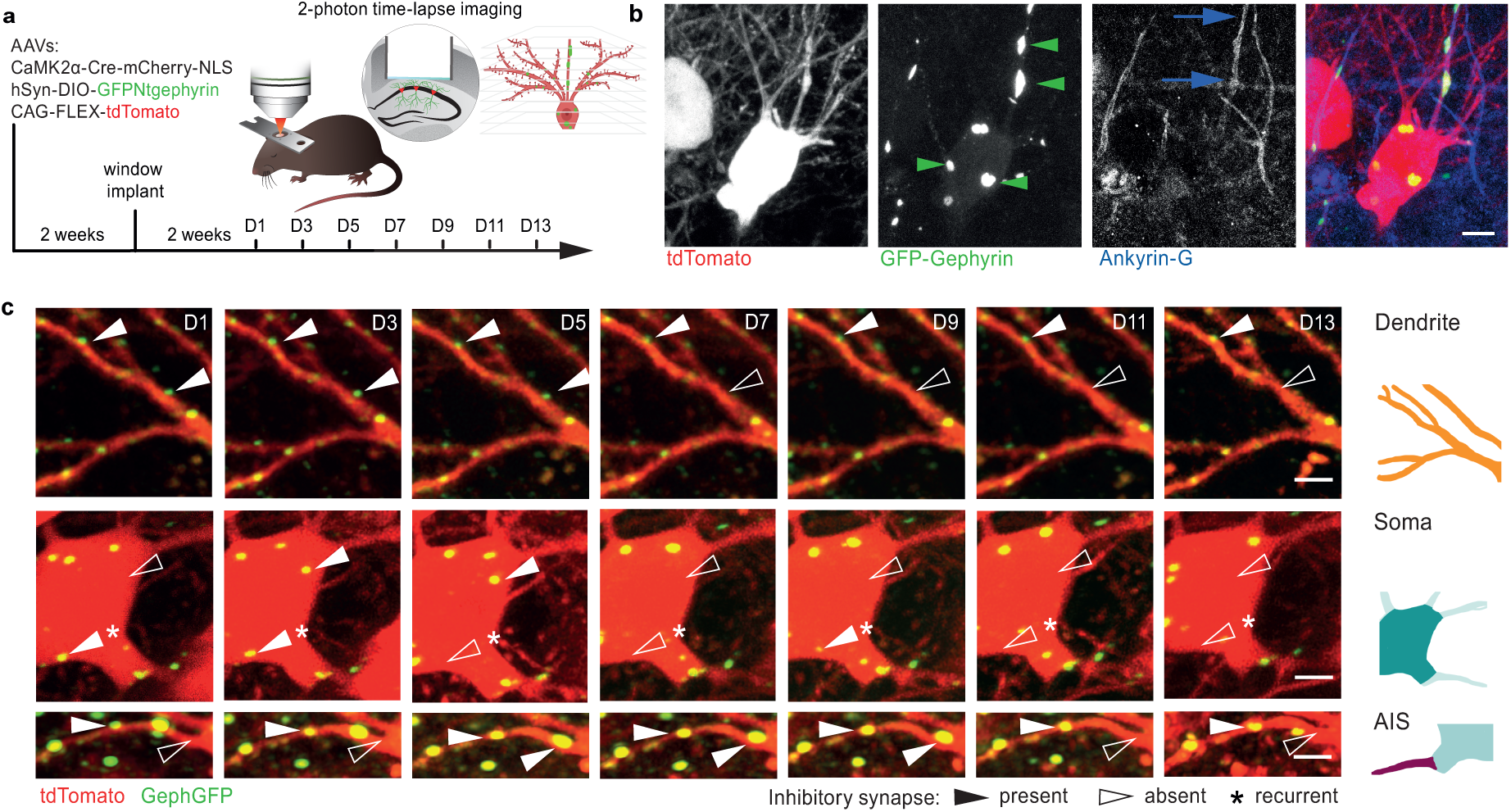
Longitudinal tracking of dCA1 PNs’ INS in different subcellular compartments. **a**, Schematic description of the strategy to label a sparse subset of dCA1 PNs and the INS impinging on them as well as of the experimental timeline. **b,** Confocal images (Maximum Intensity Projection, MIP, of 55 slices, z-step 0.25 µm) of dCA1 PNs expressing GFP-Gephyrin and tdTomato and immuno-stained for ANK-G. Green triangles indicate INS, blue arrows indicate a neurite positive for ANK-G. Scale bar, 5 μm. **c,** Two-photon time series of dCA1 PN subcellular compartments expressing GFP-Gephyrin and tomato over 13 days. Solid and empty triangles indicate present and absent synapses, respectively. Asterisks mark recurrent synapses. (MIPs of 15-40 slices, z-step 1 µm). Scale bar, 5 µm.

### Dynamics of INS area are compartmentalized on dCA1 PNs

First, we quantified the area of INS in different subcellular compartments of dCA1 PNs (**Fig. 2a**) by identifying circular regions of interest centered in the middle of Gephyrin-GFP-positive puncta and counting the number of pixels whose fluorescence was comprised within two S.D. of the mean fluorescence inside the region (**Fig. 2b**). Dendritic INS were smaller than somatic and AIS INS while AIS INS were bigger than dendritic and somatic INS (**Fig. 2c** and **Fig. S2a**). This was not surprising as AIS INS are compound cartridge synapses that contain clusters ^28^ exceeding 2P microscopy’s resolution limit. We treated separable AIS cartridges as single units and refer to them as AIS INS.

**Figure 2.**
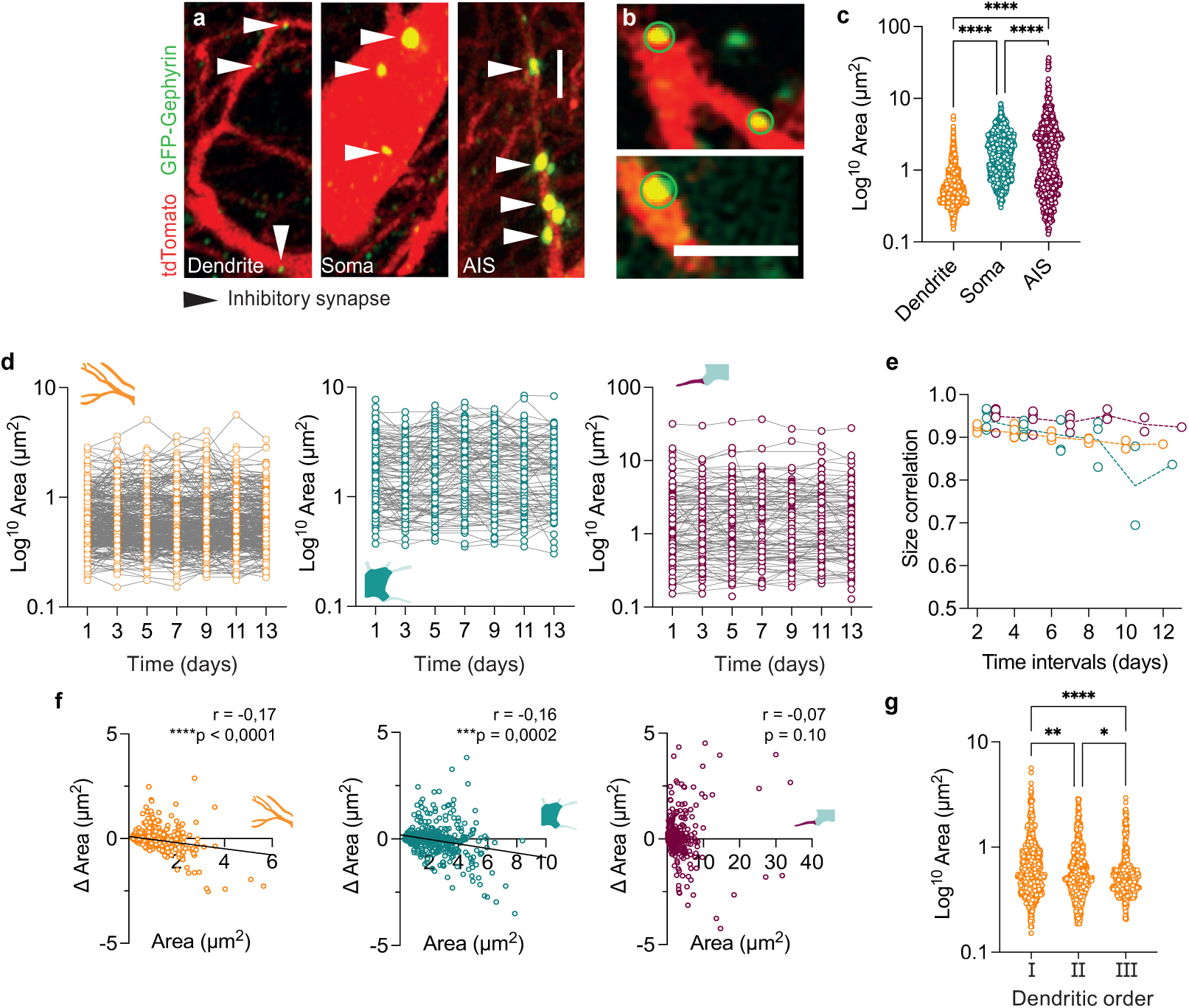
Compartmentalized properties of INS’ size. **a,** Two-photon images (MIPs, 15-40 slices, z-step 1 µm) of GFP-Gephyrin-positive INS (green) on dCA1 PNs subcellular compartments (dTomato, red). Triangles indicate synapses. Scale bar, 5 µm. **b,** Two-photon images (single Z planes) of GFP-Gephyrin-positive INS (green) on dCA1 PNs (tomato, red). Green ovals are example Regions Of Interest (ROIs) used to measure INS size. Scale bar, 5 µm. **c,** Size distributions of dendritic (orange), somatic (teal) and AIS (purple) INS. All time points pooled. **** p < 0.0001; Kruskal-Wallis test with Dunn’s correction for multiple comparisons; n_dendrite_ = 2287, n_soma_ = 1047, n_AIS_ = 743). **d,** All dendritic (orange, right), somatic (teal, middle) and AIS (purple, right) INS areas tracked over 13 days. Dendritic INS: 382 day average, 358-401 range. Soma INS: 153 day average, 136-165 range. AIS INS: 106 day average, 100-113 range. **e**, Values of significant (*p* < 0.05) Spearman correlations between areas of dendritic (orange), somatic (teal) and AIS (purple) INS at each time point versus any other time point. **f**, Correlations between the sizes of dendritic (orange, left), somatic (teal, middle) and AIS (purple, right) INS at each time point and the variation in size between that time point and the next one. All time points pooled. Values in panels refer to Spearman correlations. n_dendrite_ = 2104 pairs, n_soma_ = 537 pairs, n_AIS_ = 558 pairs. Lines represent linear fits to the data. **g**, Distributions of sizes of dendritic INS sorted by dendritic order. \**p* = 0.011, \*\**p* = 0.0023, \*\*\*\**p* < 0.0001; Kruskal-Wallis test, with Dunn’s correction for multiple comparisons; n_DO1_ = 848, n_DO2_ = 759, n_DO3_ = 674.

As INS area correlates to the number of synaptic receptors and to synaptic transmission efficacy ^33–35^, tracking the areas of hundreds of INS through time (**Fig. 2d** and **Fig. S2b**) enabled us to investigate the temporal stability of the patterns of synaptic inhibition on PNs. Analysis of the pairwise Spearman correlations between each time point and all others for INS areas (**Fig. 2d**) revealed a remarkable level of similarity through time, with correlations between neighboring time points as high as correlations between time points far apart from each other (**Fig. 2e** and **Fig. S2c**, **d**). Moreover, areas of dendritic and somatic INS on a given day correlated negatively to changes in area between that day and the following day (or delta area, Δarea) (**Fig. 2f**, orange and teal, respectively), which did not occur in AIS INS (**Fig. 2f**, purple). Finally, INS area decreased with increasing distance from the soma (**Fig. 2g**).

### Long-term dynamics of INS are compartmentalized on dCA1 PNs

Somatic INS were significantly less stable than dendritic or AIS INS, with higher 48h Gain, Loss and Turnover Fractions (**Fig. 3a**). These high rates translated to a remarkably fast decay of somatic INS and a strong difference in survival between somatic versus dendritic and AIS INS (**Fig. 3b**).

**Figure 3.**
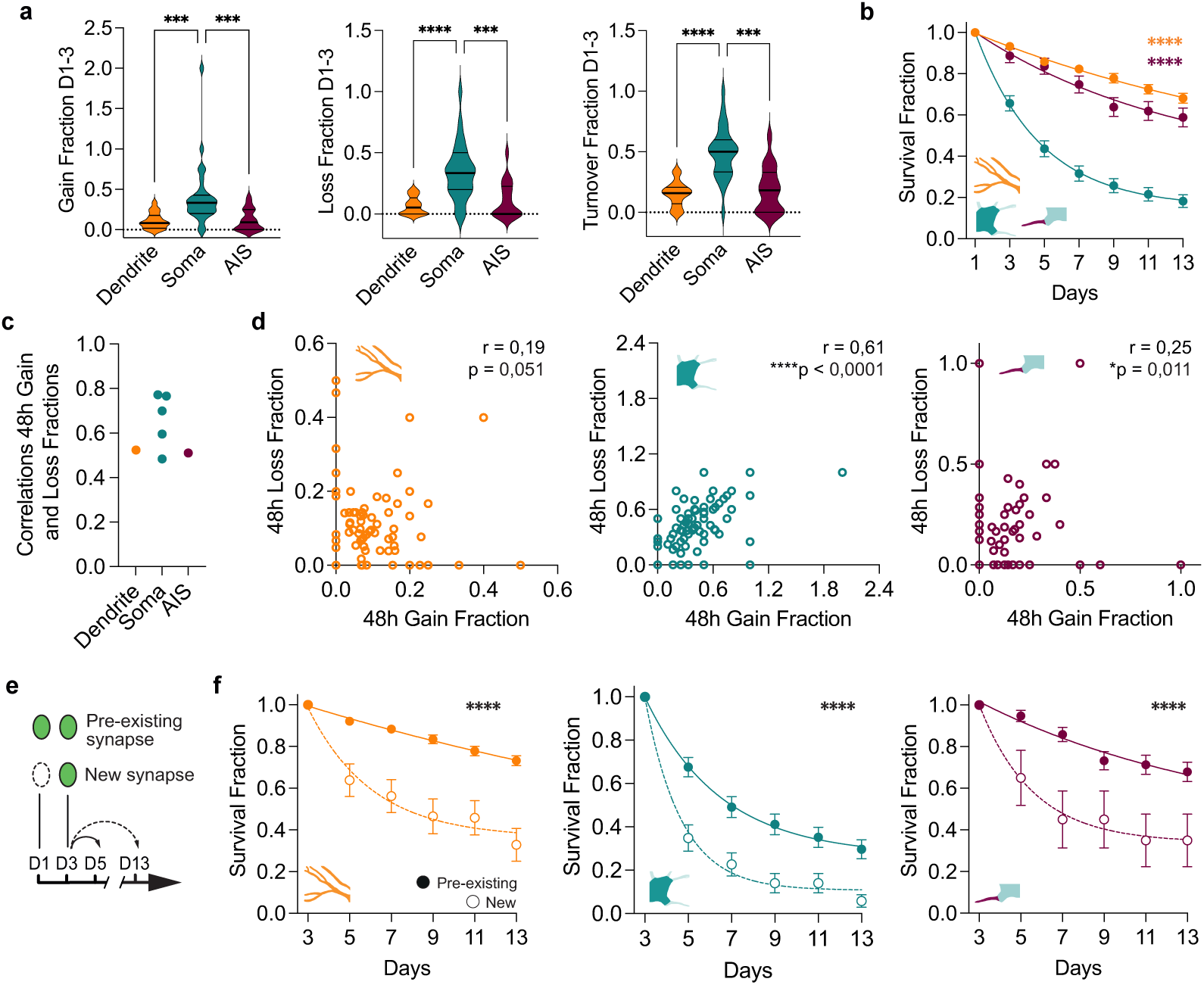
Compartmentalized properties of INS’ dynamics. **a,** Distributions of dendritic (orange), somatic (teal) and AIS (purple) INS temporal dynamics (Gain, left; Loss, middle; Turnover, right) over 48h *per* neuron. \*\*\**p* < 0.001; \*\*\*\**p*< 0.0001; Kruskal-Wallis test, with Dunn’s correction for multiple comparisons; n_Dendrite_ = 17 neurons, n_Soma_ = 27 neurons, n_AIS_ = 17 neurons. Horizontal bars are medians and quartiles of the distributions. **b**, Survival Fractions of dendritic (orange), somatic (teal) and AIS (purple) INS *per* neuron. **** *p* < 0.0001; Extra sum of squares F-Test of the fits; n_Dendrite_ = 17 neurons, n_Soma_ = 27 neurons, n_AIS_ = 17 neurons. Curves are least squares fits of one phase decay of mean (circles) and S.D. Survival Fraction values per neuron. Error bars are S.D. of bootstrapped surrogate distributions with 10^4^ iterations. **c**, Values of significant (p < 0.05) Spearman correlations between 48h Gain and Loss Fractions of dendritic (orange), somatic (teal) and AIS (purple) INS at each time point. **d**, Correlations between 48h Gain and Loss Fractions of dendritic (orange, left), somatic (teal, middle) and AIS (purple, right) INS, *per* neuron. All time points pooled. Values in panels refer to Spearman correlations. N_Dendrite_ = 102 pairs, n_\Soma_ = 162 pairs, n_AIS_ = 102 pairs. **e**, Schematic description of pre-existing and new synapses. **f**, Survival Fractions of new (empty circles) and pre-existing dendritic (orange, left), somatic (teal, middle) and AIS (purple, right) INS *per* neuron. **** *p* < 0.0001; Extra sum of squares F-Test of the fits; n_Dendrite_ = 17 neurons, n_Soma_ = 27 neurons, n_AIS_ = 31 neurons. Curves are least squares fits of one phase decay of mean (circles) and S.D. Error bars are S.D. of bootstrapped surrogate distributions with 10^4^ iterations.

As high turnover of somatic INS could result from synapses persisting but shifting their position, we wanted to measure the motion of persistent INS. We thus sampled INS as well as a series of stable landmarks as often as our technique allowed in a new set of mice (**Fig. S3a-c**) and retrieved their spatial coordinates. We then projected them in the same three-dimensional frame (**Fig. S3d** and **e**) to measure the 3D distances of each INS and landmark at each time point to itself at any other time point. There were no differences between the distances of INS and landmarks in all subcellular compartments (**Fig. S3f**), showing that the position of persistent INS is equally stable in all compartments. Thus, the higher turnover of somatic INS cannot originate from higher local fluctuations in position.

Somatic INS showed a remarkable consistency with each other at the neuronal level (**Fig. 3c**). Overall correlation between gain and loss was higher for somatic than for dendritic and AIS INS (**Fig. 3d**), but in all cases sufficient to keep the number of dendritic, somatic and AIS INS in each neuron roughly constant throughout the experiment (**Fig. S2e**).

Newborn dendritic spines are less stable than older ones, because they tend to miss functional synapses ^36–41^. Similarly, new INS (that existed for less than 48h) disappeared faster than pre-existing INS (that existed for at least 48h or longer) in all three subcellular PNs’ compartments (**Fig. 3e, f**). As our labelling strategy enables direct detection of inhibitory post-synapses, our data show that INS are also more susceptible to be eliminated early during their maturation.

### Longitudinal tracking of dendritic spines as proxies for excitatory synapses

Previous work in the visual cortex demonstrated significant differences between excitatory and inhibitory synapses’ dynamics ^22,23^, still very little is known about these differences in the hippocampus. To compare the dynamics of dCA1 inhibitory and excitatory synapses, we tracked dendritic spines - as proxies for excitatory synapses - in dCA1 PNs of Thy-GFP mice (**Fig. 4a, b** and **Fig. S4a, b**) for a duration of 13 days.

**Figure 4.**
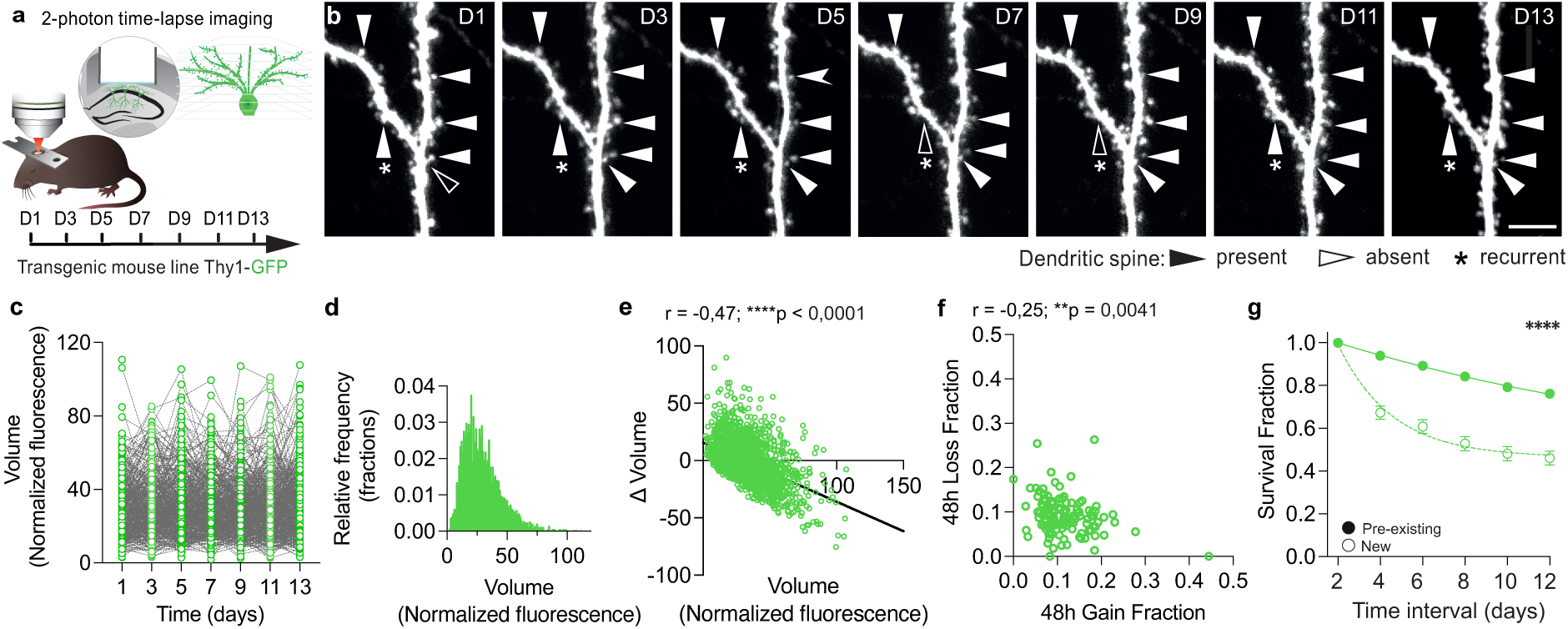
Longitudinal tracking of dCA1 PNs dendritic spines as proxies for excitatory synapses. **a**, Schematic description of the strategy to label a sparse subset of dCA1 PNs by sparse expression of somatic GFP in the Thy1-GFP transgenic mouse line as well as of the experimental timeline. **b,** Two-photon time series of dCA1 PN dendritic segments and spines expressing GFP. Solid and empty triangles indicate present and absent spines, respectively. Asterisks mark recurrent spines. Scale bar, 5 µm. **c,** Dendritic spines volumes tracked over 13 days. 464 day average, 438-529 range **d**, Distribution of spine volumes (all time points pooled, n = 3245). **e**, Correlation between the sizes of spines on each time point and the variation in size between that time point and the next one. All time points pooled. Values in panels refer to Spearman correlations. n = 2374 pairs. Line is the linear fit to the data. **f**, Correlations between 48h Gain and Loss Fractions of spines, *per* neuron. All time points pooled. Values in panels refer to Spearman correlations. n = 132 pairs. **g**, Survival Fractions of new (empty circles) and pre-existing (solid circles) spines *per* neuron. **** *p* < 0.0001; Extra sum of squares F-Test of the fits; n = 22 neurons. Curves are least squares fits of one phase decay of mean (circles) and S.D. Survival Fraction values per neuron. Error bars are S.D. of bootstrapped surrogate distributions with 10^4^ iterations.

The distribution of dendritic spines’ volumes was skewed towards higher values (**Fig. 4c, d**) - similar to dendritic INS (**Fig. S5a)** - and the volume of a spine at any given time point and Δ-Volume to the next time point were negatively correlated (**Fig. 4e**) - consistent with previous work ^42,43^-. The 48h Gain, Loss and Turnover Fractions were stable in time (**Fig. S4d**) and when we collapsed all time points, we found a small but significant negative correlation between gain and loss rates at the neuronal level (**Fig. 4f**). This however, did not lead to a measurable variability in the total number of dendritic spines over time (**Fig. S4c**). Finally, new spines disappeared much faster than pre-existing ones (**Fig. 4g**), consistent with previous work ^41^.

### Dendritic INS are less dynamic than dendritic spines

While the size distributions of dendritic INS and spines were similar (**Fig. S5a**), the size patterns of dendritic INS on each day were significantly more correlated to each other than the ones of dendritic spines (**Fig. 5a**) and, consistently, the sizes of dendritic spines fluctuated more than the sizes of dendritic INS (**Fig. 5b** and **Fig. S5b)**. This relative stability of dendritic INS could be due partially to the comparison of changes in two-dimensional area (for INS) to changes in three-dimensional volume (for spines). However, the relative stability of INS was also evident when we compared dynamics of dendritic spines and INS. Dendritic INS showed 48h Turnover Fractions significantly lower than dendritic spines (**Fig. 5c**), mostly due to smaller gain rates (**Fig. S5c**). Altogether these data show that dendritic INS are less dynamic than dendritic spines through time in dCA1 PNs.

**Figure 5.**
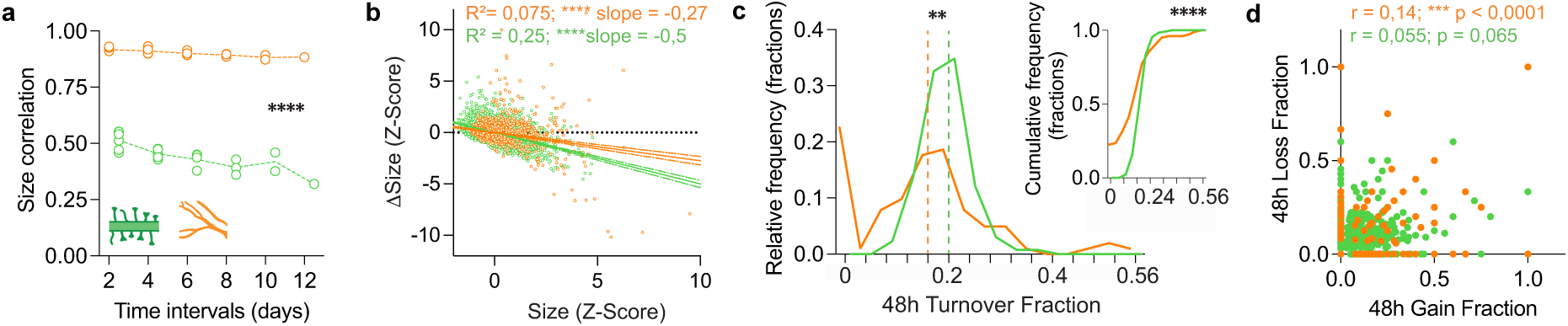
INS are more stable than dendritic spines. **a**, Values of significant (*p* < 0.05) Spearman correlations between sizes of spines (volume, green) and dendritic INS (area, orange) on each time point versus any other time point. *\*\*\*\*p* <0.0001; Mixed-effects model with Geisser-Greenhouse correction; n_Spines_ = 21 values, n_DendINS_ = 21 . **b,** Correlations between the sizes of dendritic INS (orange) or dendritic spines (green) on each time point and the variation in size between that time point and the next one. All time points pooled. Lines represent linear fits to the data and 95% confidence intervals, values in the panels refer to the fits. **c**, Distributions of 48h Turnover Fraction of spines (green) or dendritic INS (orange). \*\**p* = 0.0088; Mann-Whitney test; n_Spines_ = 132, n_DendINS_ = 102. All time points pooled. **Inset:** cumulative distributions of 48h Turnover Fraction of spines (green) or dendritic INS (orange). \*\*\*\**p* < 0.0001; Kolmogorov-Smirnoff test; n_Spines_ = 132, n_DendINS_ = 102. All time points pooled. **d**, Correlations between 48h Gain and Loss Fractions of spines (green) or dendritic INS (orange) averaged per dendritic segment. Values in the panels refer to Spearman correlation., n_Spines_ = 1098 pairs, n_DendINS_ = 744 pairs. All time points pooled.

The correlation between gain and loss rates was weak (even when significant) for dendritic INS and spines alike, when considering both the distributions at the level of the neurons (**Fig. S5d)** or dendritic segments (**Fig. 5d**). Still, the number of synapses in each dendritic segment was largely constant through time (**Fig. S5e** and **f**).

### Size and persistence correlate to different extents in dendritic spines and INS

A relationship between size and stability has been shown for dendritic spines in that bigger spines tend to survive longer than smaller spines and spines become smaller before disappearing on the time scale of hours ^44–46^. However, it is unclear whether such a relationship holds true over longer durations of time and to what extent INS also show such a relationship. Long-lived dendritic spines (observed for 7 consecutive time points, **Fig. 6a**) were significantly bigger than short-lived ones (observed for a single time point) (**Fig. 6b**, green). While the same relationship was true also for AIS INS, dendritic INS only showed a trend in this direction (**Fig. 6b**, purple and orange, respectively). Surprisingly, somatic INS showed a trend in the opposite direction with short-lived synapses being bigger than long-lived ones (**Fig. 6b**, teal). The fraction of shrinking spines was higher than the one of growing spines only for unstable dendritic spines (**Fig. 6c** and **6d**, green), while for INS there were no significant differences between shrinking and growing synapses (**Fig. 6d**, orange, teal and purple). Overall, these data show that size is a better predictor of survival for dendritic spines than for INS (with the partial exception of AIS INS) and that changes in size relevant for dendritic spines’ survival can be traced back several days prior disappearance.

**Figure 6.**
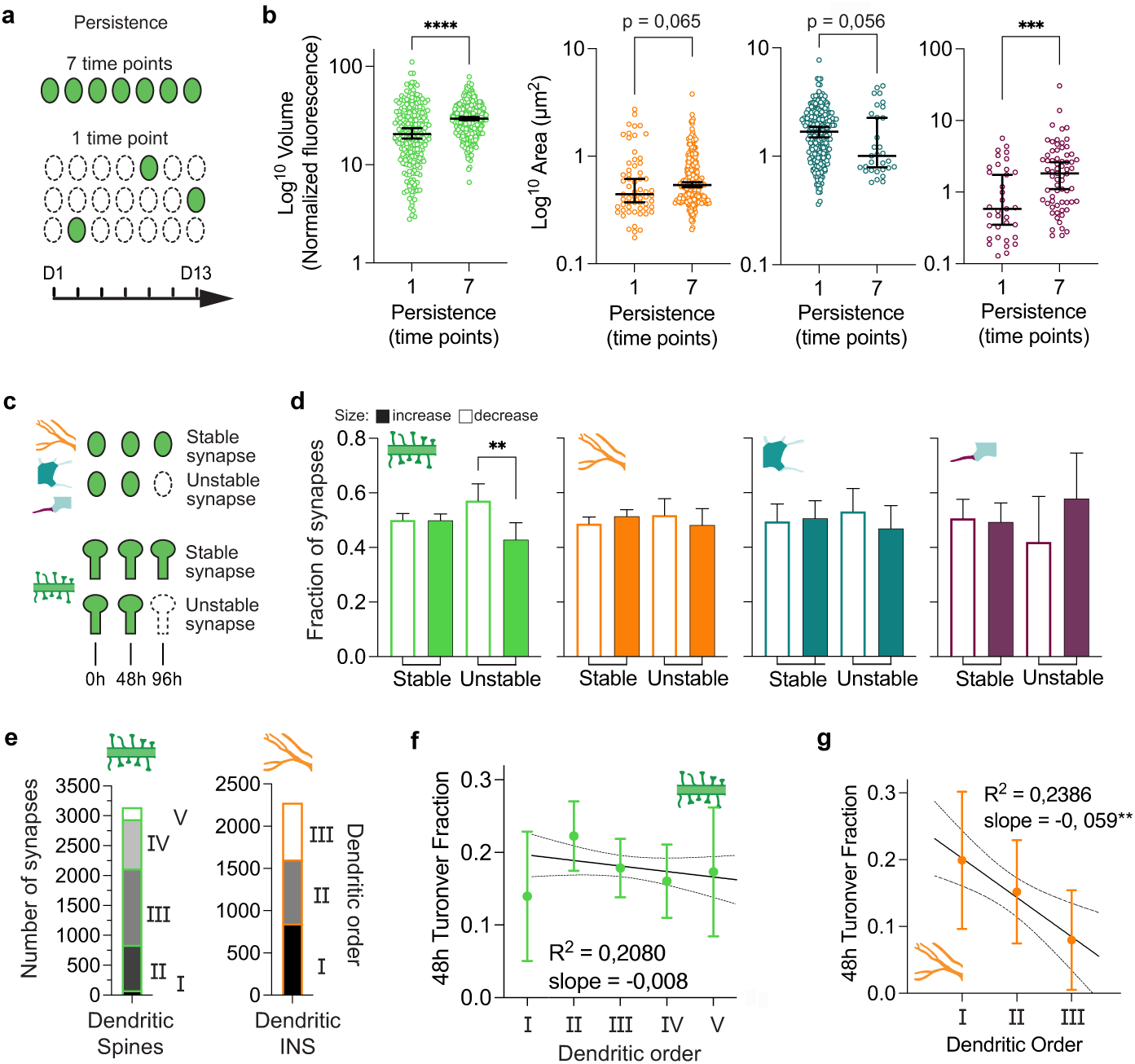
Differential relationship between size and stability amid dendritic spines and INS. **a**, Schematic description of the persistence of long-versus short-lived synapses. **b**, Distributions of sizes of spines (volume, green), dendritic (area, orange), somatic (area, teal) and AIS (area, purple) INS observed for 1 or 7 consecutive time points (\*\*\*\**p* < 0.0001, \*\*\**p* = 0.0006; Mann-Whitney test; n_1d-spines_ = 235, n_13d-Spines_ = 291; n_1d-DendINS_ = 61, n_13d-DendINS_ = 273; n_1d-SomaINS_ = 246, N_13d-SomaINS_ = 31; n_1d-AISINS_ = 35, n_13d-AISINS_ = 67). For synapses observed for 13 days are average size over 7 data points. **c**, Schematic description of stable or unstable synapses (INS, top; spines, bottom), within a three-time points sliding time window. **d**, Distributions of fractions of stable or unstable synapses (spines, green; dendritic INS, orange; somatic INS, teal; AIS INS purple) whose size decreased or increased between the first and the second time points (\*\**p* = 0.006, all others *p* > 0.39; Kruskal-Wallis test, with Dunn’s correction for multiple comparisons; n = 5 time windows). Error bars are S.E.M. **e**, proportion of dendritic spines (left, green) or dendritic INS (right, orange) sorted according to the order of the dendritic segment they were located on (roman numerals). **f, g**, Distributions of 48h Turnover Fraction of dendritic spines (f, green) or INS (g, orange) sorted by dendritic order. Values in the panel refer to the linear fit to the data. Solid line, linear fit; dashed line, 95% confidence interval of the fit. Circles are means, vertical bars are S.D.

Finally, we found a small but significant negative trend between the persistence of synapses and their distance from the cell body – measured as the order of the dendrites they are located on (**Fig. 6e**) -, with synapses located further from the soma being more stable than synapses closer to it (**Fig. 6f** and **g**).

### Learning changes inhibitory synaptic dynamics in dCA1

To investigate how learning affects structural synaptic plasticity in the hippocampus, we tracked dendritic spines as well as dendritic, somatic and AIS INS (**Fig. S6a, b)** as we subjected them to Trace Fear Conditioning (TFC) (**Fig. 7a**). TFC is a strong one-trial hippocampal-dependent learning task and mice significantly froze above baseline levels during context and tone recall (**Fig. 7b**). We calculated the 48h Gain, Loss and Turnover Fractions (**Fig. S7a - l**) and averaged the values corresponding to baseline or after TFC while leaving untouched the data points that included TFC training or Recall per each neuron (**Fig. 7c, f, i, l** and **Fig. S8a - d**). Comparisons to baseline revealed a sustained increase in turnover of dendritic INS for the whole period following TFC (**Fig. 7c**), which was accompanied by a significant decrease in long-term survival (**Fig. 7d**). At shorter time scales, stability of dendritic inhibitory synaptic patterns – *i.e.* the normalized fraction of synaptic sites that remained stable between two time points (Simple Matching Coefficient) - was significantly higher than in the shuffled dataset (**Fig. 7e**, asterisks) but TFC had no obvious effect on this. We did not detect any significant effect of TFC on somatic INS (**Fig. 7f-h**). For AIS INS, while the comparisons of Turnover Fractions with baseline did not reach significance (**Fig. 7i**), there was a significant increase in survival after TFC (**Fig. 7j**). Interestingly, this effect was also evident at the level of the connectivity patterns that became significantly more stable than the shuffle, after TFC (**Fig. 7k**, asterisks). While we detected a small decrease in turnover of dendritic spines between days 7 and 11 (**Fig. 7l**), this did not result in long-term changes in survival (**Fig. 7m**) nor differences in stability patterns of synaptic sites (**Fig. 7n**).

**Figure 7.**
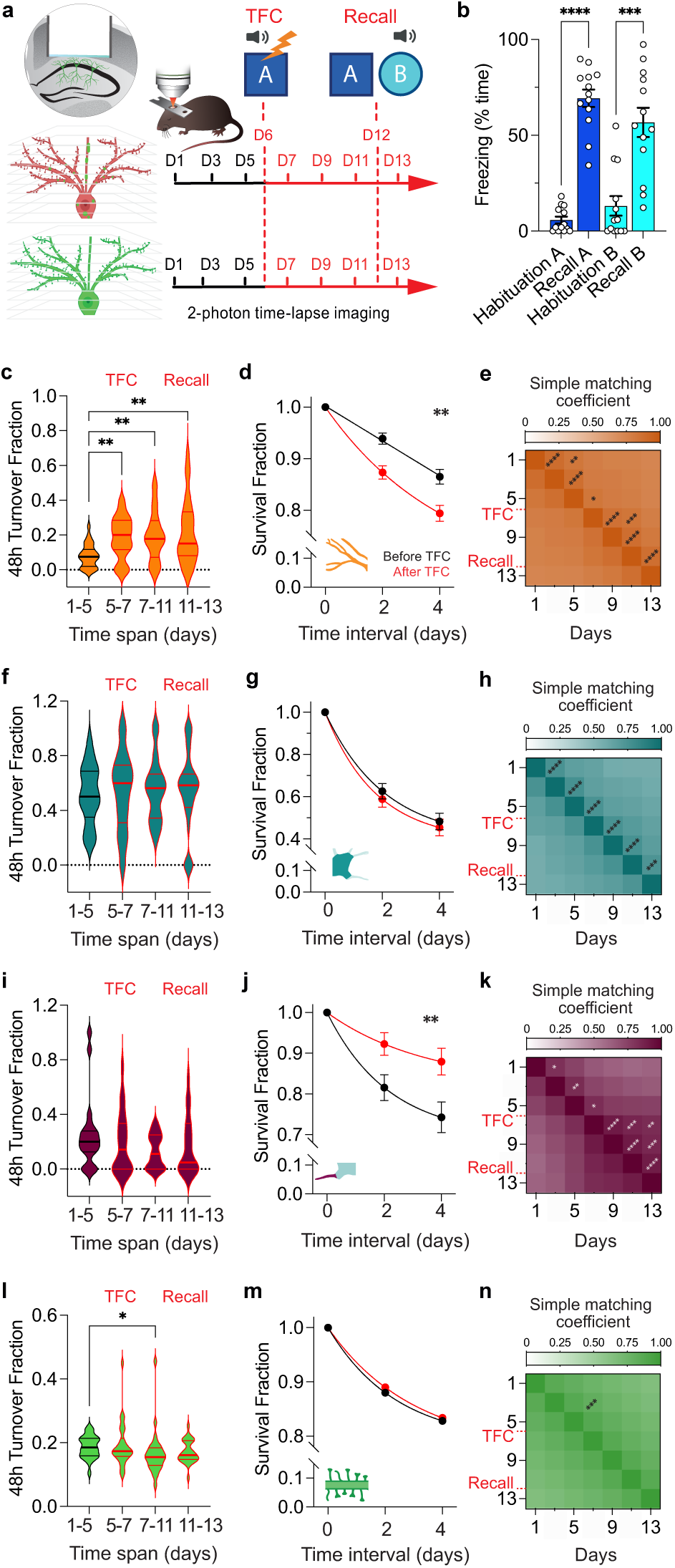
Learning a fear association changes dCA1 synaptic dynamics. **a**, Schematic description of dendritic spines and INS labelling as well as of the experimental timeline. **b**, Distributions of freezing time during habituation to Context A (blue, left), recall in Context A (blue, right), habituation to Context B (turquoise, left) and recall in Context B (turquoise, right). **** *p* < 0.0001, \*\*\**p* = 0.0007; Friedman test, with Dunn’s correction for multiple comparisons: n = 13 mice. Circles represent single mice, bars are means and error bars are S.E.M. **c, f, i, l**, Distributions of 48h Turnover Fractions (c, spines, green; f, dendritic INS, orange; i, somatic INS, teal; l, AIS INS, purple) over different epochs (prior to TFC, across TFC training, after TFC training and across recall) *per* neuron. \**p* = 0.02, \*\**p* < 0.004; Kruskal-Wallis test, with Dunn’s correction for multiple comparisons: n_spines_ = 26 neurons, n_dendAIS_ = 27 neurons, n_somaINS_ = 28 neurons, n_AISINS_ = 19 neurons. Horizontal lines are medians and quartiles of the distributions. **d, g, j, m**, Survival Fractions of spines (d), dendritic INS (g), somatic INS (j), AIS INS (m) before (black) and after TFC training (red) *per* neuron. Extra sum of squares F-Test of the fits; d, spines, *p* = 0.8265; n = 26 neurons; dendrite INS; g, \*\**p* = 0.0041; N = 27 neurons; j, soma INS\*\**p* = 0.0041; N = 28 neurons; m, AIS INS, \*\**p* = 0.00301; n = 19 neurons. Curves are least squares fits of one phase decay to the mean (circles) and S.D. Error bars are S.D. of bootstrapped surrogate distributions with 10^4^ iterations. **e, h, k, n,** Simple Matching Coefficients of binary site vectors pooled from all dendritic (e, orange) somatic (h, teal) and AIS (k, purple) INS or spines (e, green). Red dashed lines indicate the days of TFC and Recall. Permutation test \**p* < 0.05, \*\**p* < 0.01, \*\*\**p* < 0.001, \*\*\*\**p* < 0.0001 are significances for differences to shuffle data. n_Dendrite_ = 911 sites, n_Soma_ = 446 sites, n_AIS_ = 177 sites, n_Spines_ = 4753 sites.

Overall, our data shows that learning a fear association affects mostly INS, rather dendritic spines’ dynamics. Importantly, TFC led to opposite outcomes for INS located in different subcellular compartments with rearrangement of INS located on the dendrites and stabilization of INS located on the AIS of dCA1 PNs.

### Computational modelling to investigate the role of AIS-projecting neurons on the local network inhibitory connectivity patterns

AIS-projecting inhibitory neurons are ideally positioned to control the output of dCA1 PNs ^30,47^ and indeed they have been shown to control the activity of place cells during spatial navigation ^48^. We thus wanted to investigate the role of these neurons on structural INS plasticity. To this aim we generated a neural circuit model simulating the main features of the local dCA1 connectivity and focused on the changes in dynamics of structural INS plasticity occurring when we specifically changed parameters of AIS-projecting inhibitory neurons. Our model included (i) excitatory input PNs (**Fig. 8a**, blue) some only active during the training paradigm (**Fig. 8a**, pink outline), (ii) excitatory output pyramidal neurons – with distinct dendrite, soma and AIS compartments – (**Fig. 8a**, black), and 4 types of inhibitory neurons classified according to their connectivity to subcellular compartments of output neurons (**Fig. 8a**, orange for dendritic-projecting, teal for mostly somatic-projecting, purple for AIS-projecting and grey for inhibitory neuron-projecting) and whose proportions were based on literature ^9,10,49–61^. Importantly, as we also wanted to test the effect of learning on INS plasticity, we used a reinforcement learning setup that mimicked our experimental TFC protocol (**see Methods**). The simulated circuit model was used to control the behavior of an agent navigating a virtual environment. During baseline, the agent navigated the environment retrieving random positive rewards, then it was exposed to a novel environment where the stimulation of sound and shock neurons simulated the TFC protocol by delivering a negative reward. We tuned the amplitudes of the stochastic contribution to rewiring to qualitatively match the baseline turnover curves obtained experimentally during baseline (**Fig. 8b**), then we fixed the model parameters and applied the TFC protocol. The resulting INS turnover dynamics in the model following TFC were largely consistent with our experimental results, with stabilization of AIS INS and no significant changes in somatic INS dynamics (**Fig. 8c**, right and middle respectively). However, the model did not reproduce the TFC-dependent destabilization of dendritic INS (**Fig. 8c**, left).

**Figure 8.**
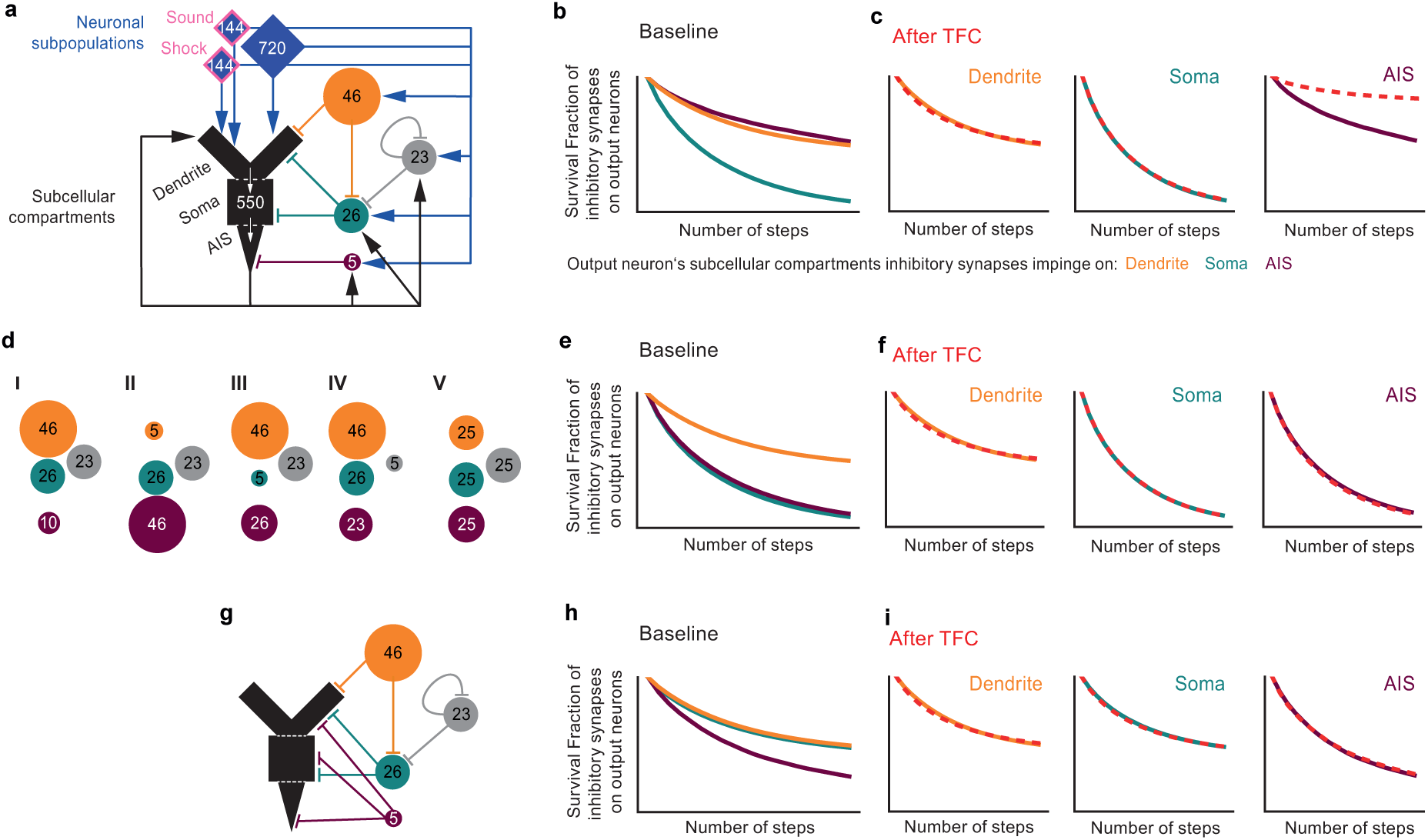
Computational modelling to investigate the role of AIS-projecting neurons on the local network inhibitory connectivity patterns. **a**, Schematic description of the model of the local neuronal network. Excitatory input (blue and blue with pink outline) and output (black) PNs with distinct dendrite, soma and AIS compartments, dendritic-projecting (orange), somatic-projecting (teal) AIS-projecting (purple) and inhibitory neuron-projecting (grey) inhibitory neurons. Numbers indicate the number of neurons implemented in the model. **b**, Survival Fractions of modelled dendritic (orange), somatic (teal) and AIS (purple) INS during the baseline period. **c**, Comparisons of the Survival Fractions of modelled dendritic (orange, left), somatic (teal, middle) and AIS (purple, right) INS during the baseline period versus after TFC (dashed red lines). **d**, Schematic description of the changes in the number of inhibitory neurons projecting to the dendritic (orange), somatic (teal), AIS (purple) compartments or to other inhibitory neurons (grey). **e**, Survival Fractions of modelled dendritic (orange), somatic (teal) and AIS (purple) INS during the baseline period after any of the changes described in d. **f**, Comparisons of the Survival Fractions of modelled dendritic (orange, left), somatic (teal, middle) and AIS (purple, right) INS during the baseline period versus after TFC (dashed red lines) after any of the changes described in d. **g**, Schematic description of the changes in the connectivity of inhibitory neurons projecting to the AIS. **h**, Survival Fractions of modelled dendritic (orange), somatic (teal) and AIS (purple) INS during the baseline period after any of the changes described in g. **i**, Comparisons of the Survival Fractions of modelled dendritic (orange, left), somatic (teal, middle) and AIS (purple, right) INS during the baseline period versus after TFC (dashed red lines) after the changes described in g.

We then explored different circuit architectures and quantified the resulting changes in INS survival, to investigate the role of AIS-targeting inhibitory neurons on INS structural dynamics. Partial removal of excitatory drive to the AIS-projecting inhibitory neurons - by removing either excitatory synapses from output neurons or excitatory synapses from input neurons (**Fig. S9a, I** and **II**) - did not have significant effects on INS connectivity dynamics during baseline or after TFC (**Fig. S9b**). Increasing the number (**Fig. 8d I**) or the proportion of AIS-projecting inhibitory neurons (**8d II - V**), led to marked instability of AIS INS (**Fig. 8e**) and prevented the stabilization of AIS INS after TFC - which was observed in the experimental *in vivo* data - (**Fig. 8f**). Changing the sign (from inhibition to excitation) of AIS INS synapses or preventing activity of AIS-projecting inhibitory neurons also led to instability of AIS INS (**Fig. S9e** and **f**). Simply decreasing the number of the neurons belonging to the disinhibitory population (**Fig. S9c**) also let to marked baseline instability of AIS INS but it did not prevent stabilization after TFC (**Fig. S9d**). Finally, we tested the effect of the connectivity pattern, rather than the activity levels, of AIS-projecting inhibitory neurons on INS stability. To this aim we set AIS-projecting inhibitory neurons to project also to all other compartments of excitatory output pyramidal neurons (**Fig. 8g**). Under these conditions AIS INS became more unstable during baseline while somatic INS became more stable (**Fig. 8h**) and exposure to TFC did not affect INS dynamics in any subcellular compartment. (**Fig. 8i**). We obtained a very similar result when we allowed all three classes on inhibitory neurons projecting to PNs to synapse on all subcellular compartments (**Fig. S9**).

Overall, our model shows that baseline stability as well as stabilization of AIS INS after TFC are a function both of the small proportion of AIS-projecting inhibitory neurons as well as of their specific pattern of connectivity.

## DISCUSSION

### Longitudinal tracking of INS in different subcellular compartments of dCA1 PNs

Investigating dynamics of excitatory synaptic structural plasticity in the brain of live animals lead to a leap in our understanding of brain plasticity and its role in learning and recall ^19–21^. By comparison inhibitory synaptic structural plasticity is understudied and previous studies tackling it in rodents mostly focused on neocortical areas ^22–27^. Here we labelled and tracked INS and dendritic spines on dCA1 PNs over two weeks during baseline and upon learning a hippocampal-dependent memory task. Tracking the persistence of INS through time and the stability of their size patterns enabled us to investigate the long-term efficacy of synaptic inhibition on PNs, as the efficacy of synaptic transmission on longer time scales is a function of the physical persistence and size of a synapse ^33–35^. Finally, we used quantitative mathematical modelling to investigate the properties of AIS-projecting inhibitory neurons that might underlie learning-induced changes in INS dynamics.

### Subcellular compartmentalization of INS size

The size of inhibitory synapses is variable and we find some of this variability to map on specific neuronal subcellular compartments with dendritic INS being smaller than somatic INS. This compartmentalization could be due to different classes on inhibitory neurons projecting selectively to the dendritic arbors and somata ^10,58,62,63^, spatial clustering of several INS on the soma or AIS ^29–31^ and reflect different strengths of synaptic transmission. The size of INS located on AIS was the largest and with the broadest distribution, as expected since AIS INS are cartridge synapses and are likely formed by unresolved clusters of smaller synapses.

All INS showed a remarkable stability in size over time thus reflecting the ability of synapses to maintain their individual properties over long time-scales (an ability often referred to as synaptic tenacity ^64^). Consistent with previous data ^42,64,65^, also in the live brain we find evidence of a mechanism regulating the size of dendritic spines as well as of INS located on the dendrites and somata of dCA1 PNs by decreasing the size of bigger and increasing the size of smaller synapses at the time scale of days. Size regulation of INS located in the AIS did not follow the same principles, probably because of the compound nature of these INS or because of their specific role in directly regulating neuronal output.

### Subcellular compartmentalization of INS dynamics

INS located in different subcellular compartments also showed different levels of stability with dendritic being the most stable and somatic being the least stable INS. This compartmentalization might stem from the specific activity patterns or genetic makeup of pre-synaptic inhibitory neurons ^10^ but also from interactions between the pre- and post-synaptic neurons ^66,67^. Segregation of INS dynamics might contribute to the soma-projecting inhibitory neurons’ ability to synchronize activity of large neural populations ^12,47,61,63,68–70^ and to the role of dendrite-projecting inhibitory neurons in controlling local dendritic nonlinear integration and plasticity processes ^11,71–73^.

Somatic INS showed a turnover significantly higher than dendritic and AIS INS. This was not due to fluctuations in the position of somatic INS and it cannot be only explained by clustering of several synapses (as also AIS INS are clustered but they do not show higher turnover - besides, clustering of synapses leads to apparent stability ^74^). The higher somatic INS turnover was surprising as it is unclear how a high level of turnover might support synchronization of the activity in large populations of PNs over long time scales. Interestingly, however, high temporal variability of somatic inhibition might support - if not underlie - constant changes in the patterns of cellular plasticity ^75^ and activity ^76–78^ which is a very prominent phenomenon in dCA1. Moreover, turnover of dendritic and somatic INS might have fundamentally different consequences on neuronal connectivity. In fact, dendritic-projecting inhibitory neurons mostly have a single synaptic contact on a postsynaptic dendritic segment ^8^ - thus loss of a single INS is slated to have a large effect on the excitability of a dendritic segment – while somatic-projecting inhibitory neurons have multiple INS on the soma of the same postsynaptic PN ^71^ - thus the loss of a single INS will have a smaller effect on the connectivity of pre- and postsynaptic partners -. This redundance of inhibitory somatic connectivity might thus render the synchronization of PNs robust in the face of high INS turnover.

### Dendritic INS are less dynamic than spines

We find INS located on basal dendrites in the Stratum Oriens of the dCA1 to be more stable than dendritic spines in the same brain area in terms of size and dynamics. While this stability seems at odd with previous work reporting an overall shorter persistence of INS in comparison to spines ^22,23^, our work differs in three main ways from previous ones, which can explain this mismatch. First, our study focuses on the hippocampus - where dynamics of excitatory synapses are significantly different from neocortical ones ^74^ – and might indicate also dynamics of INS are brain-area specific. Second, we studied basal dendrites, rather than apical dendrites as in the neocortical studies. Notably, the inhibitory connectivity between the basal and apical domain is very different, which could underlie different dynamics of INS located in apical and basal dendrites. Third, INS in the basal aspect of dCA1 predominantly target dendritic shafts ^8^. This is important as in the visual cortex most dynamic events were restricted to INS innervating dendritic spines while shaft INS were significantly more persistent ^22,23^. Hence INS located on dendritic shafts could be the most persistent synaptic population irrespective of the brain area. Overall, despite the apparent differences with previous work our data corroborates the idea that shaft INS function as relatively time-invariant regulators of global dendritic excitability (including control of the spread of back-propagating action potentials) while spine INS function as flexible controllers of local dendritic events ^22,23^.

In the future, it will be interesting to track INS and dendritic spines in the same neurons to be able to investigate the local relationship between these two classes of synapses to higher level of detail.

### The relationship between size and persistency in dendritic spines and INS

Dendritic spines persisting for 13 days were larger than ones observed a single time. This shows that in the dCA1 larger spines – that generally contain larger synapses – survive longer also when analyzed at longer time scales. Also dendritic INS showed a trend in the same direction, indicating that also larger INS – with larger, mature synapses ^7,34^ – tend to survive longer. Somatic INS showed a trend going in the opposite direction. However, the number of long-lived somatic INS was small - due to their high turnover - which precludes reaching a definitive conclusion. Finally, also the AIS INS persisting for 13 days were larger than the ones observed a single time, thus indicating that even for cartridge synapses the total synaptic area is a predictor of synaptic stability.

Dendritic spines became smaller within 48 hours before disappearing while INS did not. Dynamic changes in dendritic spine’s size occur within a few hours upon artificial triggering *ex-vivo* ^79,80^. Size decrease 48h before disappearance shows that dendritic spines can be slated for elimination already days before they actually disappear. By comparison, the selection of INS to eliminate seems to have faster time scales as there is no significant INS shrinking 48h before disappearing. This is consistent with the remarkable stability of INS size. Altogether these data show that that the dynamic range of sizes that each spine explores during its lifetime is bigger than the one of INS.

Interestingly, for both dendritic spines and INS, dynamics decreased along the proximal-distal axis in basal dCA1 dendrites. Increased structural persistence could compensate for electrotonical distance from the soma as more persistent physical connection between pre- and postsynaptic partners should enable a more stable synaptic connection. This compensation – together with the regenerative properties of local ion channels ^81,82^, increasing number of synaptic receptors with distance from the soma ^83^ and pushing the balance between excitatory and inhibitory synapses in favor of the former ^61^ – could balance distance-dependent attenuation of distal synaptic inputs in basal dCA1 dendrites.

### Learning significantly affects INS’ dynamics in dCA1

Previous work showed dynamics of INS in the visual cortex upon sensory deprivation ^22–24,26^ and in the motor cortex upon motor learning ^25^, we now demonstrate learning-related changes in INS in the dorsal aspect of the hippocampus, a brain area important for episodic learning.

Turnover of basal dendritic INS increased immediately after TFC and remained high for several days, leading to their destabilization. This is consistent with previous work in the motor cortex ^25^, and indicates that long-term decrease in dendritic inhibition in conjunction with restructuring of the pattern of inhibition in the basal dendrites of dCA1 PNs supports learning of a fear association. Changes in the AIS INS were of opposite sign to dendrites with a significant stabilization of the overall AIS inhibitory connectivity pattern upon TFC. Inhibition at the AIS can very effectively prevent the generation of action potentials ^84–87^, thus the stabilization of AIS INS connectivity pattern should lead to stabilization of the PNs output pattern. Therefore, while on the one hand TFC leads to rearrangements in dendritic inhibition - which likely change dendritic computations in single neurons and in turn the activity patterns of dCA1 PCs -, on the other hand it leads to stabilization of the PCs output patterns. Overall, differential control of dendritic and AIS inhibition by structural INS plasticity has the potential to effectively regulate the long-term changes in activity patterns of large populations of PNs.

The fact that dynamics of somatic INS did not respond to TFC was unexpected, as the number axonal boutons of parvalbumin-expressing inhibitory neurons - which mainly inhibit the soma and peri somatic regions of PNs - increase in the motor cortex upon training ^25^. A possible explanation of this mismatch might lay in the different kinetics of learning used in the two studies. In fact, Chen and colleagues used a behavioral paradigm that lasted several days and the increase in boutons they detected was gradual, over several days. This would suggest that single-shot learning such as TFC does not elicit significant structural somatic INS plasticity.

Finally, while learning a fear association affected the dynamics of INS, it did not significantly impact dendritic spines in dCA1 PNs. Previous studies demonstrated plasticity and learning-induced changes in spines dynamics in the neocortex ^41,46,88–95^ ^96–98^, but no previous work addressed the effect of fear learning on spine dynamics in the hippocampus. Moreover, changes in dendritic spines’ density after fear conditioning have been reported in the past in the hippocampal CA1 ^99,100^, but these results were based on comparison of different groups of mice and the magnitude of these changes was small (0,2 to 0,5 spines/um increase in density in basal dendrites). Since, the mice we imaged clearly learned the fear association, our data show that learning a fear association affects the spines’ dynamics in dCA1 to a much lower extent than in other brain areas and possibly to such a limited extent to prevent detection by 2P microscopy.

Further experiments, should clarify whether different learning paradigms, including ones based on positive feedback or with longer time scales, have different effects on structural hippocampal synaptic dynamics.

### Computational modelling suggests properties of AIS-projecting inhibitory neurons underlying the learning-induced changes in INS structural connectivity patterns

We established a neural circuit model simulating the main features of single PNs and of local dCA1 connectivity, and reproducing the main features of *in vivo* INS dynamics. We then used this model to systematically investigate INS structural dynamics in the main subcellular compartments of PNs during baseline and upon learning.

Stabilization of AIS INS upon TFC – a prominent feature of the *in vivo* dataset – was robust across a variety of experimental conditions and parameters. We identified three basic local circuit features leading to stabilization of AIS synapses. The first is sparsity of connectivity, as stabilization upon TFC only occurred when the number of AIS INS synapses was small. The second is patterned connectivity, as stabilization upon TFC only occurred when AIS INS synapses originated from a single interneuron class and when this class only projected to the AIS. The third is compartmentalization of connectivity, as changing the proportions of interneurons and thereby the specific local distribution of inhibitory synapses along the compartments drastically diminished baseline stability and prevented TFC-induced stabilization of AIS INS. Interestingly, AIS INS stabilization upon TFC was not or minimally dependent on the activity rates of AIS-projecting inhibitory neurons (as decreasing excitatory input to these neurons did not prevent TFC-induced stabilization) but it was dependent on the activity of AIS-projecting inhibitory neurons to be inhibitory, as flipping the sign of AIS-projecting inhibitory neurons’ activity abolished TFC-induced AIS INS stabilization.

Overall, our mathematical modelling suggests that the lower abundance together with the specific connectivity support the key role of AIS-projecting inhibitory neurons during learning-induced stabilization of AIS inhibitory synaptic patterns.

Theoretical and experimental work has shown how the presence of a dendrites and structured connectivity can improve computational aspects of biological and artificial neuronal networks ^101–104^. In the future, it will be important to test how patterned structural synaptic turnover on hippocampal dendrites might underlie the function of the hippocampus in learning and spatial navigation.

## METHODS

### Animals

Adult female and male, 8-12 week old C57BL6/N mice and heterozygous Thy1-eGFP [Jackson laboratories Tg(Thy1-EGFP)MJrs] mice were used for INS and dendritic spine imaging, respectively. Mice had ad libitum access to water and food and were group-housed with a maximum of 5 mice per cage. Mice were held on a 12/12 hours light/dark cycle in temperature and humidity-controlled holding rooms. Experiments were conducted during the 12-hours light period. Mice were randomly assigned to experimental or control groups before the start of each experiment and each cohort consisted of both control and experimental mice. All animal procedures conformed to Max Planck Society and Leibniz Association guidelines, and they were approved in the License of animal experimentation (42502-2-1738 LIN).

### Surgical procedures

Stereotaxic injections in the dCA1 (anteroposterior (AP) = -2.30 mm, mediolateral (ML) = 1.80 mm, dorsoventral (DV) = −1.40 mm) were performed according to standard procedures 2 weeks prior hippocampal window implantation. To label inhibitory post synapses with GFP and pyramidal neurons with tomato, we injected intracranially a mixture of three Adeno Associated Viruses: AAV8-CamK2⍺-Cre-mCherry (UNC Vector Core, 3.5×10^12^ vg/mL), AAV2/1-hSyn-DIO, GFP-Geph (Vector Biolabs, 3.6×10^12^ gc/mL) and AAV1/2-CAG-FLEX-tdTomato (MPI in house production, 3.5×10^12^ vg/mL) at a volume ratio of 24:6:1 (titer ratio: 14.4:3.9:5.8). The plasmid for GFP-Geph AAV production was a kind gift of Dr. Pablo Mendez, Cajal institute, Madrid, Spain. Hippocampal window implantations were performed according to standard procedures ^105^ and local regulations. 2P *in vivo* imaging started after a minimum of 2 weeks recovery.

### Trace Fear Conditioning

On the day of training (D6), mice were placed into a conditioning chamber (19 cm x 19 cm, metal grid floor, white light illumination, and ethanol odor (Context A) in a sound isolated box (Panlab, Barcelona, Spain). After 3 min of habituation, mice received 4 mild electric foot shocks (0.75 mA, 1 s duration, US) paired to a tone (9 kHz, 20 s duration, CS) with a trace of 20 s between the tone and the shock and an interatrial interval of 105 s. On D12, mice were placed into Context A for 3 min and later, mice were put into a novel Context B (ø 15 cm diameter round transparent chamber with bedding, white light illumination, and acetic acid odor). In this new Context B, the CS tone was played for 1 min. The position of the mouse was tracked, and the freezing behavior was monitored and quantified with ANY-maze software (Stoelting, Wood Dale, IL, USA). The freezing response was calculated as the percentage of time during which the mice were not moving over the total time of probing the memory.

### Tissue preparation and immunohistochemistry

Mice were intracardially perfused with ice-cold 1x phosphate-buffered saline (PBS), followed by 4% paraformaldehyde (PFA) in PBS. Next, brains were collected and post-fixed in 4% PFA-PBS for 24h at 4°C and then transferred to 30% sucrose in PBS for 48h at 4°C. Brain slices (40 μm thick) were cut with a vibratome (Thermo Fisher Scientific, Microm or Leica) and stored in 1x PBS containing 0.025 % Sodium Azide. Prior to staining slices were first incubated in 150 mM glycine in PBS for 15-30 minutes at room temperature, followed by three 5 min-washes in PBS. Slices were then incubated in blocking buffer (10% normal goat serum, 0.8 % Triton X-100 in PBS) for 1-2 hours at room temperature, followed by three 5 min-washes in PBS. Slices were then incubated with AnkyrinG primary antibody (chicken polyclonal, Synaptic Systems 386006) diluted 1:500 in antibody buffer (5% goat serum, 0.4 % Triton X-100 in PBS) at 4°C with gentle shaking for 22 hours, followed by three 5 min-washes in PBS. Slices were then incubated in secondary anti-chicken Alexa 405 antibody (goat polyclonal, Abcam ab175675) diluted 1:500 in antibody buffer at 4°C with gentle shaking for 22 hours, followed by three 10 min-washes in PBS After staining, slices were mounted on Super Frost plus glass slides (Thermo) and allowed to dry protected from light. Once dry, the samples were covered with either Mowiol or Vectashield Hard Set mounting medium, and a cover glass was placed on top. The slides were left to dry for at least 4 hours or overnight under cover, and were subsequently stored at 4°C.

### Confocal Imaging

Confocal Images were acquired using a Leica TCS SP8 X confocal microscope equipped with a Leica HC PL APO 63x/1.40 OIL CS2 oil-immersion objective (2048 x 2048 pixel sampling, 0.0225359 x 0.0225359 x 0.2498629 µm^3^/voxel)

### In vivo imaging

Longitudinal 2P imaging was performed in anesthetized mice during the light phase every 48 h for 13 days. Each mouse underwent a previous imaging session to acquire overview images and determine appropriate laser power levels per image which were then maintained through time (range of 30-60 mW). Animals were placed on a 37°C heating pad for the duration of each imaging session and eyes were covered with Bepanthen eye ointment. Each mouse head was manually aligned to the light path as previously described ^105,106^ prior to each imaging session. Mice were image using either a Bruker Ultima IV microscope - equipped with an InSight Deep See dual color laser system (Spectra-Physics), a 1.0 NA, 25x water immersion objective (Olympus XLPlan N 25x/1.00 SVMP) and a galvanometric scanner - or a Thorlabs Bergamo microscope – equipped with a Chameleon Ti:Sapphire dual color laser system (Coherent), a 0.8 NA 40x water immersion objective (Nikon NIR Apo 40x/0.80W DIC N2) and a resonant scanner-. TdTomato and GFP were co-excited using 1040 nm and 920 nm wavelengths and their fluorescence signals were split in two in the emission path using filters and detected with by fluorescence was detected by two GaAsPs (Hamamatsu or Thorlabs 2102/M). 3D stacks were acquired at 512 x 512 pixel width (0.094 x 0.094 x 1 µm^3^/voxel size, 50-100 z-planes, 20 repetitions per each z-plane at the Bruker system or 0.1079 x 0.1079 x 1 µm^3^/voxel size, 50-100 z-planes,15 repetitions per each z-plane at the Thorlabs system). Filter sets used to split the specifically distinguish the two channels were MDF-GFP (Thorlabs), et750sp-2p_t560lpxr-UF1, t570lpxr-UF1(Chroma).

### Processing of 2P image stacks

First each z-plane that was repeatedly acquired was registered to the mean of all repetitions of that z-plane with a Rigid Body registration and then averaged to a single plane. For 2 channel images the red channel (cytosolic tdTomato) was used for reference and the same transformations were then applied to the green channel. Next, all registered and averaged z-planes were concatenated to a 3D image stack for deconvolution. 3D blind deconvolution was performed using AutoQuantX3 software and 10 iterations per stack. We then semi-automatically equalized the z-dimension in size and section of all time point 3D image stacks to concatenate them into an ImageJ timeseries 4D stack. To correct for motion artifacts in time we used a registration based on cross correlation. For 2 channel images, the red channel was used as registration reference.

### Tracking of synapses and their size over time

Prior to synaptic tracking the experimenter was blinded to the experimental groups. To track inhibitory synapses and spines on 4D timeseries image stacks we used the ImageJ Plugin, TRACKMATE ^107^. Each synapse was detected at the plane where it had its maximum diameter and it was assigned a unique identifier (ID) that included x, y and z positional coordinates for the synapse. When a synapse disappeared and re-appeared at what appeared to be the same location, we assigned a new ID.

To calculate the area of INS, we defined a circular region of interest (ROI) for each identifier and adjusted the size of the ROI to roughly match the size of the visible GFP signal. We then Z-scored all pixel intensities within the ROI and selected pixel intensities within 2 Z-scores from the mean intensity of all pixels belonging to the INS. Finaly, we multiplied this value for the pixel size to obtain the INS area.

To calculate the areas of dendritic spines, we segmented single dendritic spines and then matched these segmented spines to the tracking data. Tracked locations further refined using iterative closest point based metric ^108^. To segment dendritic spines, we implemented a convolutional neural network (CNN) trained on a large-scale dendritic spine dataset from a previous study ^109^ and fine-tuned with additional annotations from this *in-vivo* dataset. This allowed the network to generalize across different imaging conditions and accurately segment spines and dendrites in both *in-vitro* and *in-vivo* datasets. The segmentation involved extracting localized image patches centered on the manually tracked spine locations, which were then passed through the CNN. The network output consisted of a probability map highlighting spines and dendrites, which was thresholded and converted into a binary segmentation mask. To further refine the segmentation and enhance dendritic visibility, we applied a fibermetric-based enhancement filter optimized for bright, elongated structures corresponding to dendrites and spines. This method is based on a Hessian-derived multiscale Frangi vesselness filter. The fibermetric function does not classify structures but instead enhances their visibility, making them more distinguishable from the background, which is particularly useful for dendrites and spines, as it highlights their tubular morphology while suppressing non-relevant background features. This was followed by anisotropic diffusion filtering, which reduced noise while preserving fine morphological features. After contrast adjustment, a ROI - including both the dendritic shaft and the spine head - was extracted for each spine. We measured spine volume through the integration of fluorescence intensity within the segmented spine regions, with local background subtraction applied to improve accuracy. The CNN-generated segmentation was assigned a confidence score based on classification probability, providing a reliability measure for each detected spine. Overlay images were generated to display the segmented spines on the original images, allowing for direct comparison between tracked and segmented spines. Temporal changes in spine morphology were visualized across time points, and the final dataset included spine volume, area, dendritic fluorescence intensity, and segmentation confidence scores, all of which were exported for statistical analysis.

### Synaptic dynamic measures and statistical analysis

*48h Gain Fraction* was calculated as the number of synapses born (N_gained_) on each imaging day (D_n_) divided by the number of synapses present on the previous day (D_n-1_). *48h Loss Fraction* was calculated as the number of synapses lost (N_lost_) on D_n_ divided by the number of synapses present on D_n-1_. 48h *Turnover Fraction* was as the sum of (N_gained_ + N_lost_ on D_n_ divided by the sum of synapses at D_n_ and D_n-1_. Survival Fraction at each time point was calculated as the number of synapses surviving from the first to any other day (N_surviving_) divided by the number of synapses on day1 (D_1_). *Simple Matching Coefficient* between binary vector pairs of synaptic patterns at different time points was calculated as the number of synaptic sites that did not change (i.e. that were either occupied or not occupied on both days) divided by the total number of sites. For statistical comparisons we generated a permuted SMC matrix with shuffled row values (sites) while preserving the number of 1s in each column (days), thereby ensuring the number of synapses per day remained constant while the occupancy pattern of synaptic sites was shuffled. P-values were then calculated by comparing the original SMC matrix to the distribution of permuted matrices. Number of permutations was 10^4^.

*Intrasynaptic Euclidean distances* between the same synapses in consecutive time points in 3D space, were calculated using the following formula:

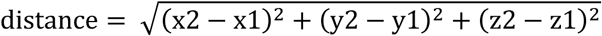

#### Permutation Test

As first step, the observed test statistic is computed: delta mean values per neuron over the 13 day time span of synaptic Survival Fractions compared in a pairwise manner across compartments. Then the Survival Fractions from all neurons and the compartments to compare are pooled. A total number of 10^4^ permutations are performed. For each permutation, the merged data is shuffled, and two new groups are formed by randomly splitting the shuffled data into two sets according to the sizes of the original groups. The test statistic (delta mean) between the two groups for each permutation is calculated and recorded. To determine the p-value, the proportion of permuted test statistics that are equal to or exceed the observed test statistic is calculated. For multiple comparison correction in comparisons of all 3 compartments, Bonferroni correction was applied, by multiplying the p-value it by the number of tests.

#### Bootstrapping

All site matrices of all neurons are pooled into one matrix. The size of each bootstrap sample equals the number of matrices in the pooled matrix (bootsize). A bootstrap sample is drawn by randomly sampling matrices from the pooled matrix with replacement until the bootstrap sample size equals the bootsize. This procedure is repeated iteratively 10^4^ times to generate the surrogate distribution where each bootstrap sample represents one resampled dataset.

We used non-parametric statistical test corrected for multiple comparison when distributions to be tested were non-Gaussian.

### Modelling and simulations

All simulations were carried out by using the CoBeL-RL simulation framework using the Tensorflow machine learning library ^110^. The simulations consisted of a virtual agent, navigating a virtual environment. The virtual agent was controlled by an artificial recurrent neural network implementing the hippocampal microcircuit model. The artificial agent received visual feedback in the form of naturalistic images generated using Blender 3D computer graphics software (version 2.79). The virtual environment was a squared 2 by 2 meters box, the walls of which were structured with photo-realistic textures. A point light source with adjustable color was placed above the center of the environment box. For the standard, baseline environment, the box was lit with white light and it contained a grid of 4 x 4 possible agent positions. The agent could freely move between these positions by generating actions. Images were captured at the agent’s current position using wide angle camera objects, giving a field of view of ca. 240°. Inputs corresponded to the rendered camera images scaled to 12 x 20 pixels (with 3 color channels each), giving a total of 12 x 20 x 3 = 720 visual input channels. Inputs were presented to the network by setting the activations of the 720 visual input neurons to the current values of the input channels.

#### Artificial recurrent neural network implementing the hippocampal microcircuit model

The network consisted of 1008 excitatory input, 550 excitatory output, and 100 inhibitory neurons. Excitatory input neurons were divided in three subpopulations: 720 visual, 144 sound and 144 shock neurons. Inputs were presented to the network by setting the activations of the neurons to the current values of the input channels. Sound and shock neurons were only active during TFC (see later) otherwise they were silent throughout the experiment. All excitatory input neurons projected to the dendritic compartment of all excitatory output neurons. Excitatory output neurons consisted of 3 compartments (dendrite, soma, AIS). Each compartment received input projections from a distinct set of inputs (excitatory and/or inhibitory, see Fig. 8a) and a Rectified Linear Unit (ReLU) non-linearity was applied to generate the compartment output. The 550 outputs of the axon compartment projected to (i) 4 neurons that controlled actions of the 4 different actions of the agent to move within the maze (up, down, left, right) (ii) to inhibitory neurons that synapsed on the axon initial segments of the output neurons (iii) to a set of inhibitory neurons that inhibited soma-projecting inhibitory neurons and (iv) recurrently to their own dendritic compartment. Inhibitory neurons belonged to 4 subpopulations with connectivity and relative abundance designed to recapitulate the local connectivity of the mouse dCA1 hippocampus. Inhibitory neurons were single compartment neuron models with ReLU activation to produce the output. Excitatory neurons consisted of 3 compartments (dendrite, soma, axon). Each compartment had a distinct set of input neurons and a ReLU non-linearity applied to generate the output. All inhibitory neurons and excitatory compartments used adaptive bias weights that were included in the training process. Synaptic weights were constrained to being either excitatory (positive) or inhibitory (negative). To prevent synapses from switching signs and to model synaptic rewiring, we adopted the method from ^111^. Briefly, for every synapse *i* we introduced a synaptic parameter *θ_i_* which was projected using a non-linear mapping *w_i_* = exp(*T\**(*θ_i_* – *θ*_0_)) > 0, where we chose the temperature parameter T=0.25 and offset *θ*_0_=1. Synapses with *θ_i_* ≤ 0 were interpreted as disconnected synapses and their effect on the post-synaptic neuron was set to zero. A single synapse was allowed between any pair of neurons in the circuit to simplify the simulation.

#### Simulation of the agent and TFC

The agent was trained using the deep Q-learning algorithm. Briefly, the hippocampal network model was trained to approximate the action-value function, that represents the expected time-discounted reward for given state / action pairs. We used a discount factor of 0.8. Weight updates were superimposed by random Gaussian noise with amplitude eta (η) to facilitate synaptic motility. The discount factor depreciates the value of rewards far in the future and thus encourages fast goal seeking. States were not presented explicitly (e.g. as locations in the maze), but only implicitly through visual, sound and shock inputs. One of 16 positions in a standard environment (context A) was randomly selected as reward location. Whenever the agent visited the reward location, a reward of +10 was delivered to the agent and the reward as immediately relocated to a new random position. The agent resided for 4500 time steps in context A. Connectivity dynamics were recorded during this period and labelled as Baseline. The agent was then placed in a novel context for 450 time steps. In this context, no reward was provided to the agent. After an initial delay of 90 time steps, the first TFC trial was initiated. Every TFC trial had a total duration of 90 time steps. The sound neurons were activated for 10 time steps, followed by a delay of 10 time steps. Then the shock neurons were activated for 1 time step, accompanied by a strong negative reward of -100, irrespective of the agent’s current location. After the shock a delay of 69 time steps, where shock and sound neurons were inactive, was provided. TFC trials were repeated 4 times before the agent was transferred back to context A. The activation values of sound and shock neurons were set to 1 during TFC trials, but they were silent throughout the rest of the experiment.

#### Analysis of the circuit re-wiring

We recorded synaptic parameter values in intervals of 90 time steps throughout the whole experiment to evaluate INS turnover and the rewiring of the circuit. Synapses were labelled as active if their synaptic parameter *θ_i_* was > 0 and as inactive else. The calculation of the surviving fraction was performed as for *in vivo* experiments. Noise amplitudes *η* were used to calibrate the behavior of the circuit, *i.e.* to qualitatively match the *in vivo* behavior for the baseline condition (*η =* 0.01 for dendrite, *η =* 0.2 for soma and *η =* 0.08 for AIS INS. After calibrating to reproduce baseline behavior, noise amplitudes were kept fixed during the whole experiment and for further evaluations, including the TFC analysis.

## ACKNOWLEDGEMENTS

The plasmid we used to generate the GFP-Gephyrin virus was a kind donation of Dr. Pablo Mendez (Cajal Institute, Madrid, Spain). Dr. Hongbo Jia supported us in the use of the 2P system, Dr. Judith Kaufhold and Albin Varga together with their team provided excellent care of experimental animals, Dr Wilko Altrock and his team genotyped all transgenic animals, Dr Liudmila Sosulina, Dr. Hiroshi Kaneko, Silvia Vieweg and Janina Juhe were always ready to help us on all sorts of different issues.

## AUTHOR’S CONTRIBUTION

K.H. designed and performed the experiments, analyzed the data, wrote the manuscript; K.D. performed the modelling work; Ö.A. wrote the software to calculate spine’s volume; U.F.A., M.B., S.S. and H.R.E. performed or supported the experiments; R.S. provided funding and access to instrumentation; A.A. designed the experiemtns, supervised the work, analyzed the data, wrote the manuscript, procured funding. All authors commented on the manuscript.

