## Supplementatry Figures and Captions for "Learning reorganizes dendritic and stabilizes axonal initial segment inhibitory synapses in CA1 pyramidal neurons"

### SUPPLEMENTARY FIGURES AND CAPTIONS

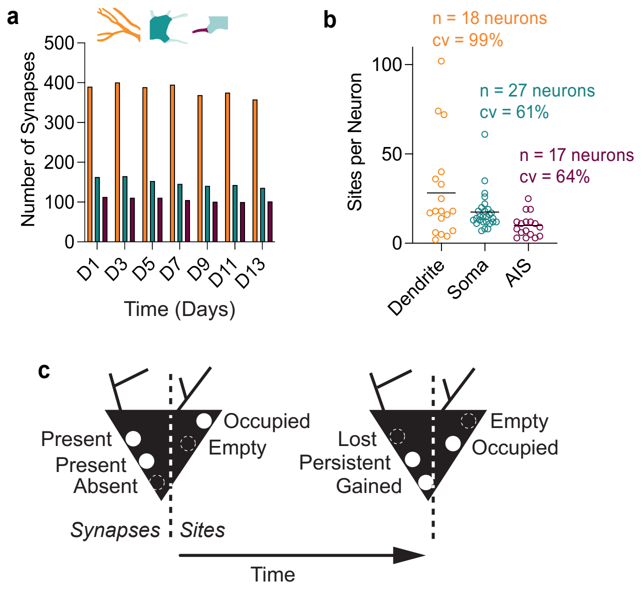


**Figure S1 | Number of INS and synaptic sites.**

**a**, Number of dendritic (orange), somatic (teal) and AIS (purple) (382, 150 and 106, average per day; 18, 27, 17 neurons respectively, 6 mice) INS per imaging day during baseline.

**b**, Number of dendritic (orange), somatic (teal) and AIS (purple) inhibitory synaptic sites per neuron during baseline.

**c**, Schematic depiction of synapses and sites on pyramidal neuron in two consecutive timepoints.

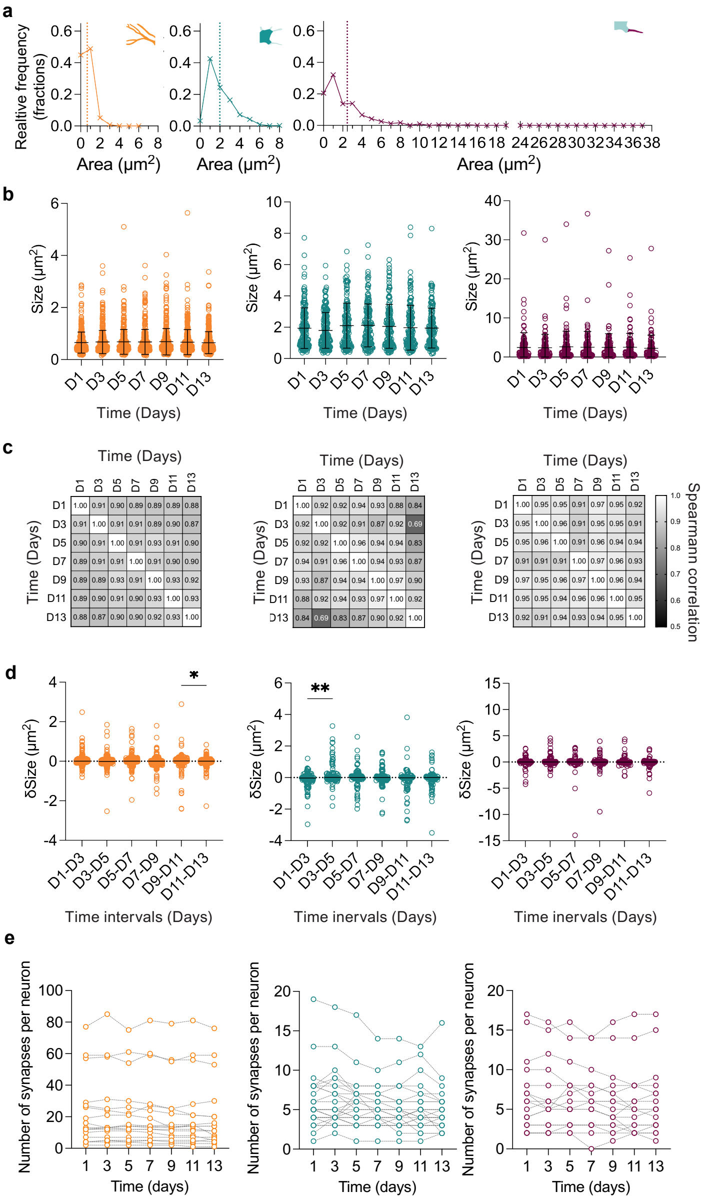


**Figure S2 | Sizes and counts of INS through time during baseline.**

**a**, Histograms of relative frequencies of synaptic sizes in each neuronal compartment, bin size 1, dotted line is mean synaptic size.

**b,** Sizes of dendritic (orange), somatic (teal) and AIS (purple) INS over 13 days. Kruskal-Wallis test, Dunn's test, $n_{Dendrite} = 358\text{-}401$; $n_{Soma} = 136\text{-}165$; $n_{AIS} = 100\text{-}113$). $p > 0.05$; Line: mean; Error bars: S.D.

**c,** Spearman Correlations matrices of sizes of INS from dendritic, somatic and AIS compartment.

**d,** Delta- sizes of dendritic (orange), somatic (teal) and AIS (purple) INS over 13 days. (Kruskal-Wallis test, Dunn's test, $n_{Dendrite} = 358\text{-}401$; $n_{Soma} = 136\text{-}165$; $n_{AIS} = 100\text{-}113$). *$p < 0.05$, **$p < 0.01$. Line: mean; Error bars: S.D.

**e,** Numbers of dendritic (orange), somatic (teal) and AIS (purple) INS *per* imaging day *per* neuron during baseline.

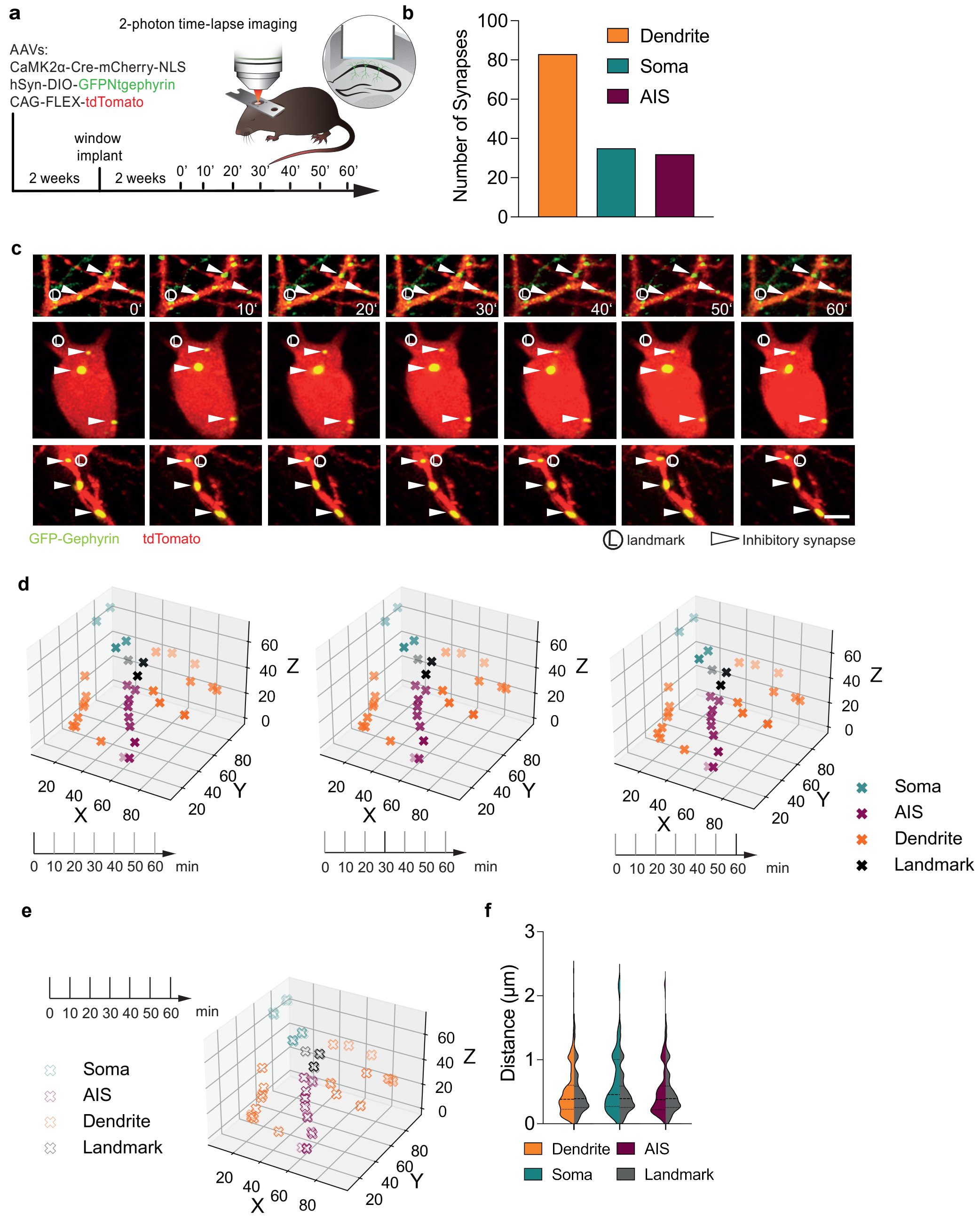


**Figure S3 | Three-dimensional euclidean distances of stable INS at short time intervals**

**a**, Schematic description of the strategy to label a sparse subset of dCA1 PNs and the INS impinging on them as well as of the experimental timeline.

**b,** Numbers of stable dendritic, somatic, AIS inhibitory sites (83, 5 and 32 respectively).

**c,** Two-photon time series of dCA1 PN subcellular compartments (dendrites, top; somata, middle; axon initial segments, bottom) expressing GFP-Gephyrin and dTomato. Solid and empty triangles indicate synapses, circulated L indicates structural landmarks. Scale bar, 5 µm.

**d,** Centroid coordinates of dendritic (orange), somatic (teal) and AIS (purple) synapses and 3 structural landmarks (black) of example neuron plotted in 3D vector space of 3 timepoints.

**e,** Overlay of dendritic (orange), somatic (teal) and AIS (purple) synapse centroid coordinates and 3 landmarks (black) of example neuron plotted in 3D vector space of 6 timepoints.

**f,** Distributions of euclidean 3D distances between adjacent timepoints of the same synapse centroid or landmark. (Landmark vs. Dendrite $p$ >0.9999*;* Landmark vs. Soma $p$ = 0.1287; Landmark vs. AIS $p$ > 0.9999; Kruskal-Wallis test, with Dunn's correction for multiple comparisons, $n_{Dendrite}$ = 470 ; $n_{Soma}$ = 167 ; $n_{AIS}$ = 66 distances)

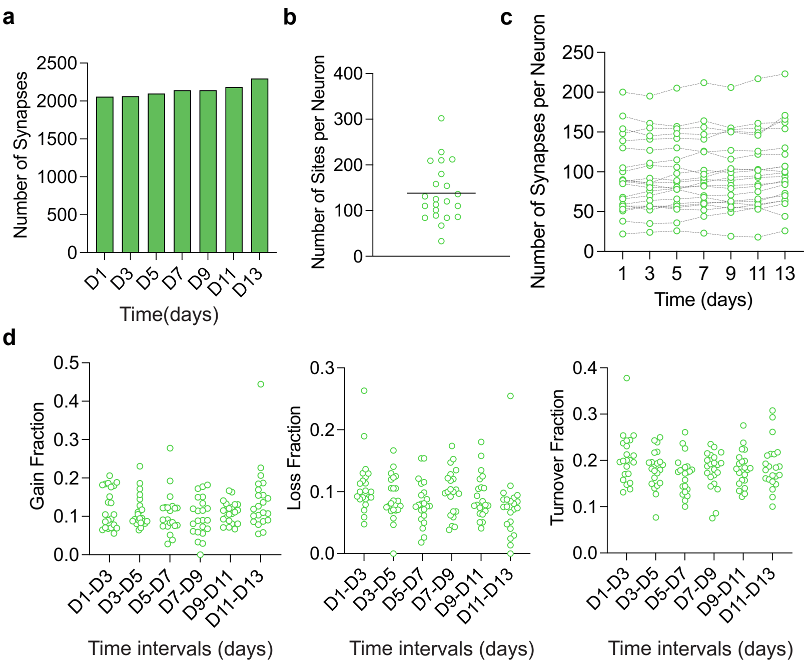


**Figure S4 | Counts of dendritic spines through time during baseline.**

**a**, Numbers of dendritic spines (2141, average per day, 22 neurons, 4 mice) *per* imaging day during baseline.

**b**, Number of dendritic spines synaptic sites (3042) per neuron during baseline.

**c**, Number of dendritic spines *per* imaging day *per* neuron during baseline. Kruskal-Wallis test, with Dunn's correction for multiple comparisons; n = 22 neurons.

**d**, Temporal dynamics of dendritic spines over 48 h (Gain, left; Loss, middle; Turnover, right). Kruskal-Wallis test, with Dunn's correction for multiple comparisons; n = 22 neurons.

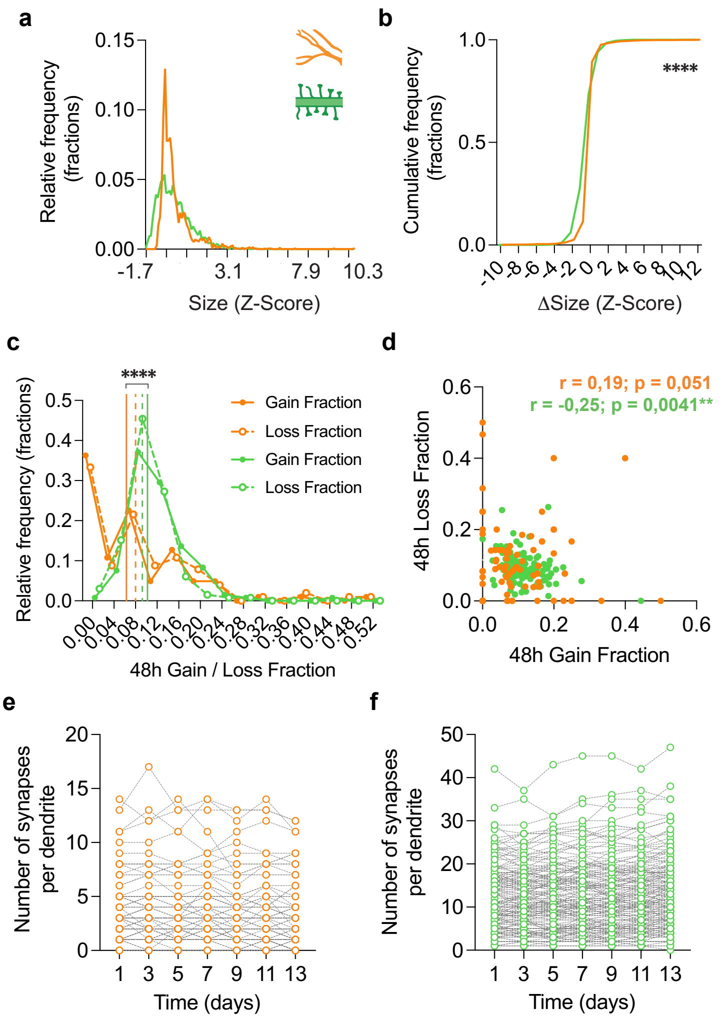


**Figure S5 | Comparison between size and dynamic properties of dendritic spines and INS.**

**a**, Histograms of Z-Scores of synaptic sizes (INS orange, dendritic spines green, $n_{Spines}$ = 3245, $n_{DendriteINS}$ = 2287, bin size 0.1).

**b**, Cumulative frequencies of Z-Scores of synaptic size deltas (INS orange, dendritic spines green, ****$p < 0.0001$, Kolmogorov-Smirnov Test; $n_{Spines}$ = 2374, $n_{DendriteINS}$ = 2104, bin size 1).

**c**, Histograms of 48h Loss (dashed line) and Gain (solid line) Fractions per neuron of INS (orange) and dendritic spines (green). ****$p < 0.0001$, Kruskal-Wallis test with Dunn's correction for multiple comparisons: $n_{Spines}$= 132, $n_{DendriteINS}$ = 102, bin size 0.04. Vertical lines are means.

**d,** Correlations between 48 h Gain and Loss Fractions of dendritic INS (orange) and dendritic spines (green) *per* neuron. All time points pooled. Values in panels refer to Spearman correlations. $n_{Spines}$ = 132 pairs, $n_{DendriteINS}$ = 102 pairs.

**e,** Number of INS *per* imaging day *per* dendrite during baseline. Kruskal-Wallis test, with Dunn's correction for multiple comparisons; n = 138 segments of 18 neurons.

**f,** Number of dendritic spines detected *per* imaging day *per* dendrite during baseline. Kruskal-Wallis test, with Dunn's correction for multiple comparisons; n = 183 segments of 22 neurons.

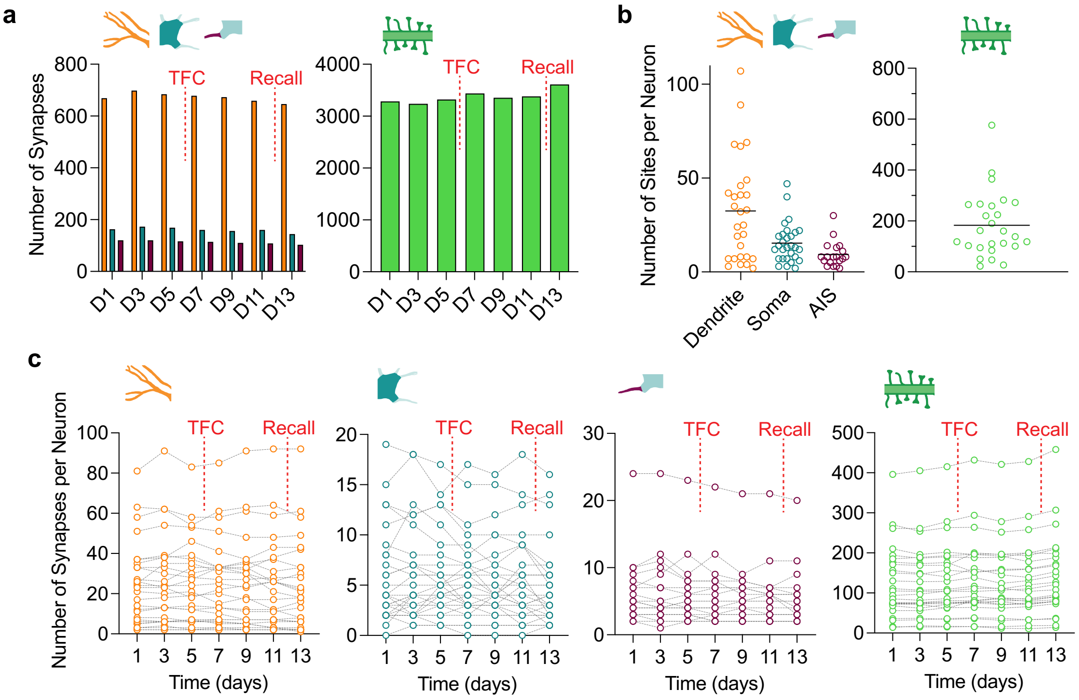


**Figure S6 | Counts of dendritic spines and INS through time in TFC group.**

**a**, Numbers of dendritic (orange), somatic (teal) and AIS (purple) INS (left) and dendritic spines (green, right) *per* imaging day, TFC group (672 dendritic, 161 somatic, 113 AIS INS and 3376 dendritic spines average per day, from 28, 29, 19, 26 neurons, 8 and 5 mice respectively). Red dashed lines indicate day of TFC training and recall.

**b**, Number of dendritic (911, orange), somatic (446, teal) and AIS (177, purple) inhibitory (left) and excitatory (4753, green, right) synaptic sites per neuron, TFC group.

**c,** Number of dendritic (orange), somatic (teal) and AIS (purple) INS and dendritic spines per imaging day per neuron, TFC group. Kruskal-Wallis test, with Dunn's correction for multiple comparisons; $n_{DendriteINS}$ = 28 neurons; $n_{SomaINS}$ = 29 neurons; $n_{AISINS}$ = 19 neurons; $n_{Spines}$ = 26 neurons.

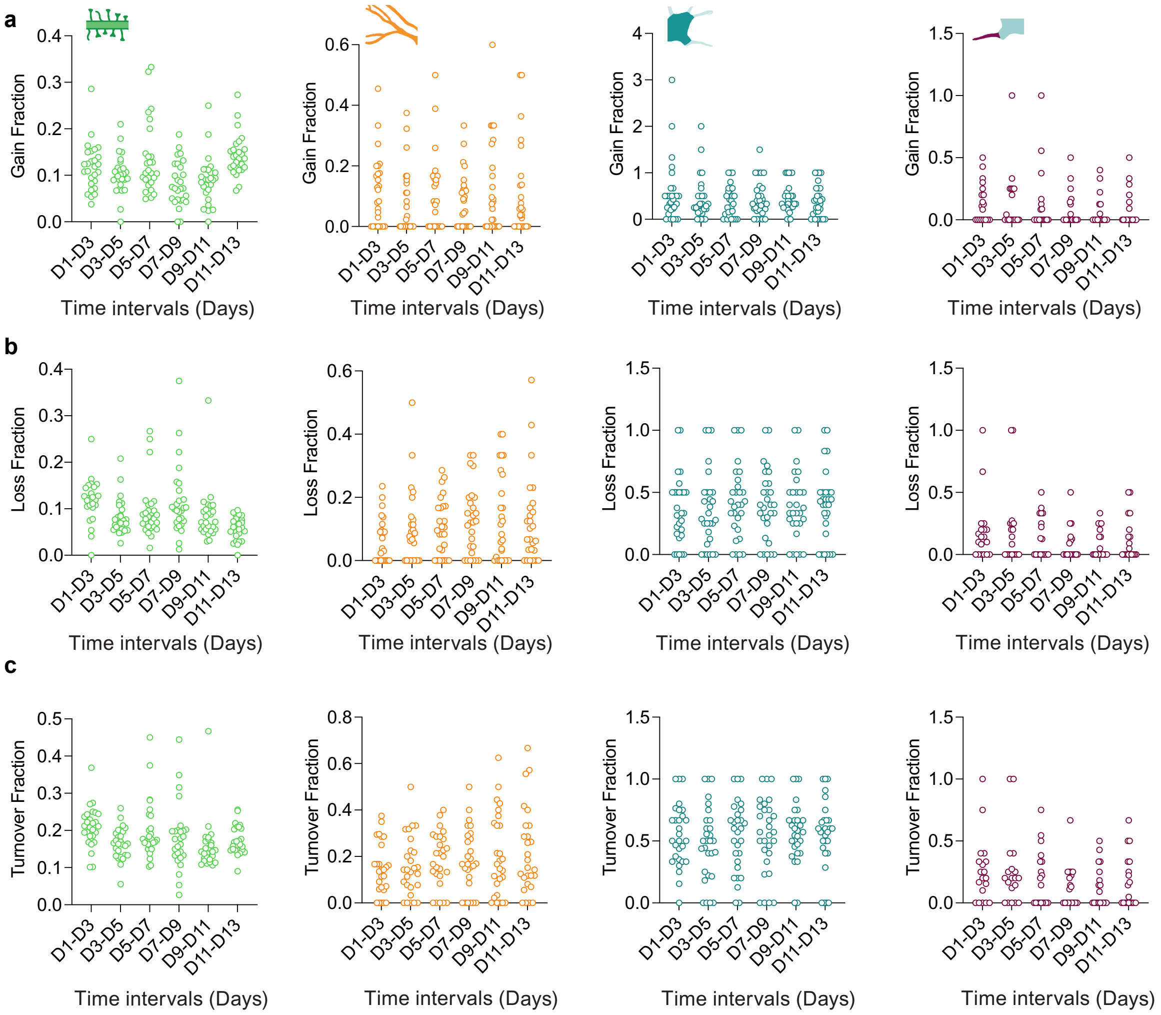


**Figure S7 | 48h Gain, Loss and Turnover Fractions of dendritic spines and INS in TFC group.**

**a - c**, Distributions of synaptic 48h Gain Fractions (a), 48h Loss Fractions (b) or 48h Turnover Fractions (c) of spines (green), dendritic INS (orange), somatic INS (teal) or AIS INS (purple). Kruskal-Wallis test pairwise day to day intervals, with Dunn's correction for multiple comparisons: $n_{Spines} = 26$ neurons, $n_{DendriteINS} = 27$ neurons, $n_{SomaINS} = 28$ neurons, $n_{AISINS} = 19$ neurons.

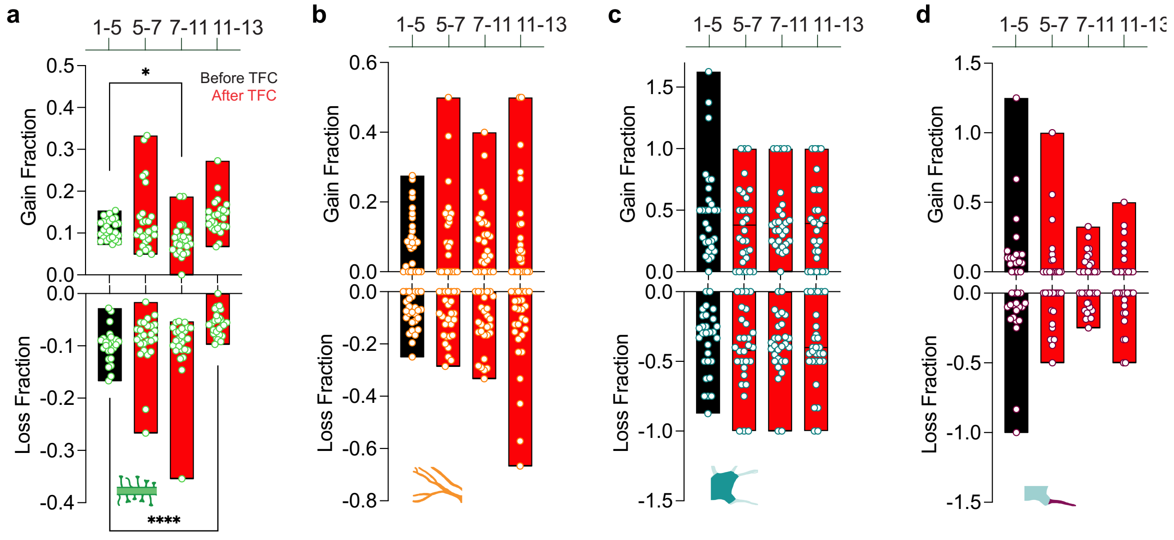


**Figure S8 | Gain and Loss of dendritic spines and INS upon learning.**

**a - d**, Distributions of synaptic 48h Gain and Loss Fractions for dendritic spines (a), dendritic INS (b), somatic INS (c) and AIS INS (d) averaged over different epochs (prior to TFC (black D1-5), across TFC training (red D5-7), post TFC training (red D7-11) and across recall (red D11-13). *$p = 0.028$, ****$p < 0.0001$; Kruskal-Wallis test each epoch to baseline, with Dunn's correction for multiple comparisons: : $n_{Spines} = 26$ neurons, $n_{DendriteINS} = 27$ neurons, $n_{somaINS} = 28$ neurons, $n_{AISINS} = 19$ neurons. Bars are data range.

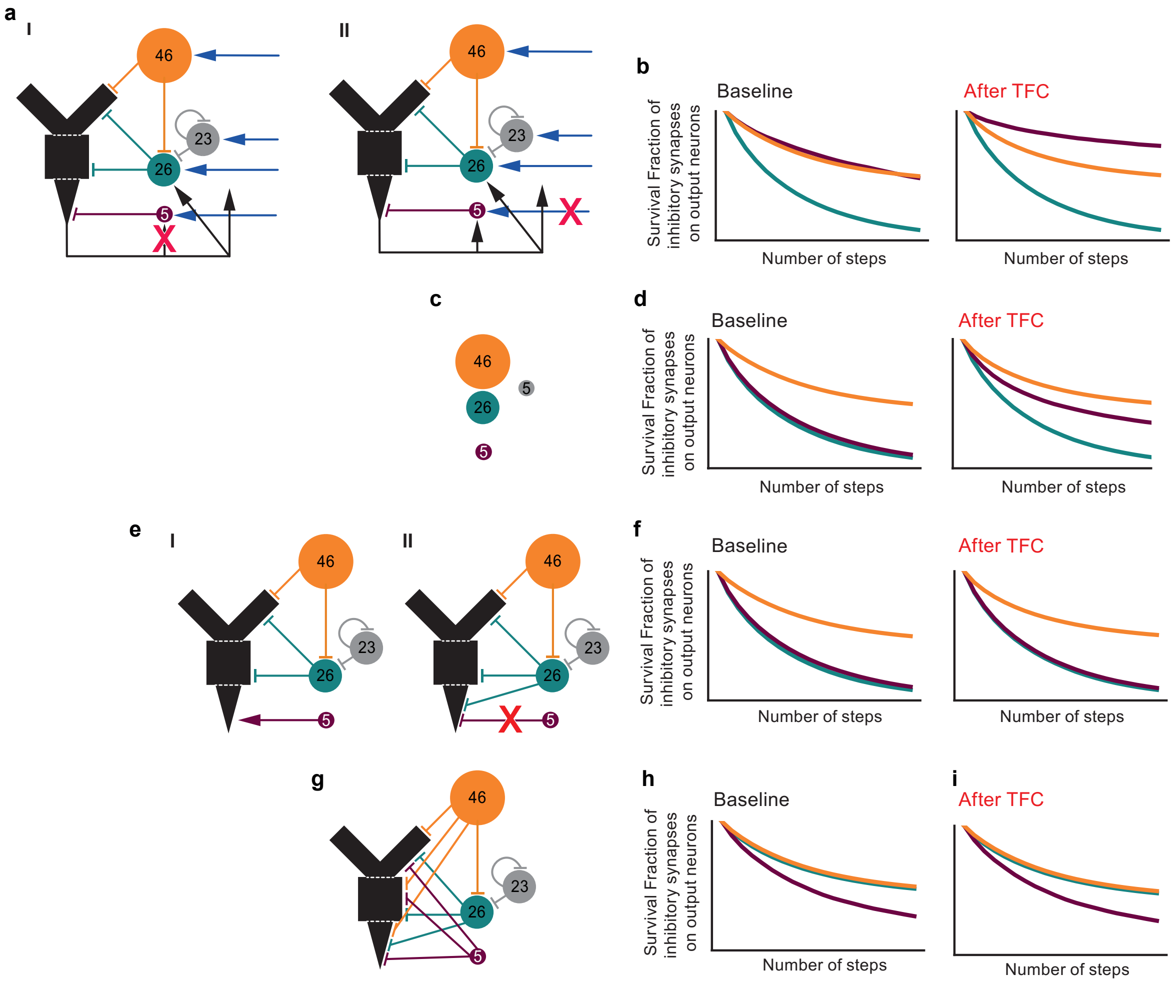


**Figure S9 | Computational modelling to investigate the role of AIS-projecting neurons on the local network inhibitory connectivity patterns.**

**a**, Schematic description of the changes in the model circuitry, crosses indicate eliminated connections.

**b**, Comparisons of the Survival Fractions of modelled dendritic (orange), somatic (teal) and AIS (purple) INS during baseline (left) versus after TFC (right) after any of the changes described in a.

**c**, Schematic description of the changes in the number of inhibitory neurons projecting to the dendritic (orange), somatic (teal), AIS (purple) compartments or to other inhibitory neurons (grey) .

**d**, Comparisons of the Survival Fractions of modelled dendritic (orange), somatic (teal) and AIS (purple) INS during baseline (left) versus after TFC (right) after the changes described in c.

**e**, Schematic description of the changes in the model circuitry, crosses indicate eliminated connections.

**f**, Comparisons of the Survival Fractions of modelled dendritic (orange), somatic (teal) and AIS (purple) INS during baseline (left) versus after TFC (right) after any of the changes described in e.

**g**, Schematic description of the changes in the model circuitry to "all to all" connectivity.

**h**, Comparisons of the Survival Fractions of modelled dendritic (orange), somatic (teal) and AIS (purple) INS during baseline (left) versus after TFC (right) after the changes described in g.
